# *χ*-separation: Magnetic susceptibility source separation toward iron and myelin mapping in the brain

**DOI:** 10.1101/2020.11.07.363796

**Authors:** Hyeong-Geol Shin, Jingu Lee, Young Hyun Yun, Seong Ho Yoo, Jinhee Jang, Se-Hong Oh, Yoonho Nam, Sehoon Jung, Sunhye Kim, Masaki Fukunaga, Woojun Kim, Hyung Jin Choi, Jongho Lee

## Abstract

Obtaining a histological fingerprint from the *in-vivo* brain has been a long-standing target of magnetic resonance imaging (MRI). In particular, non-invasive imaging of iron and myelin, which are involved in normal brain functions and are histopathological hallmarks in neurodegenerative diseases, has practical utilities in neuroscience and medicine. Here, we propose a biophysical model that describes the individual contribution of paramagnetic (e.g., iron) and diamagnetic (e.g., myelin) susceptibility sources to the frequency shift and transverse relaxation of MRI signals. Using this model, we develop a method, *χ*-separation, that generates the voxel-wise distributions of the two sources. The method is validated using computer simulation and phantom experiments, and applied to *ex-vivo* and *in-vivo* brains. The results delineate the well-known histological features of iron and myelin in the specimen, healthy volunteers, and multiple sclerosis patients. This new technology may serve as a practical tool for exploring the microstructural information of the brain.

## 1. Introduction

In neuroscience, histology has provided rich information of microscopic brain anatomy, assisting to discover neuron (Ramón y Cajal, 1893) and parcellate cerebral cortex (Brodmann, 1909). Even today, we rely on histology in exploring microscale pathogenesis of neurological diseases (Singh et al., 2004) or identifying cortical myeloarchitecture and subcortical nuclei (Ding et al., 2016). Despite the utility of histology, it inevitably entails a tissue invasion, limiting the opportunity of applying the technology to the *in-vivo* brain.

Magnetic resonance imaging (MRI) is a powerful non-invasive tool for visualizing brain anatomy and function. Various image contrasts of MRI provide the physical information of tissues such as T_1_ relaxation, T_2_ relaxation, and diffusion. Recently, efforts have been made to build connections between MRI measurements and microstructural properties of the brain (e.g., neurite density and myelination) that are not directly available at an MRI resolution (MacKay and Laule, 2016; Weiskopf et al., 2015; Zhang et al., 2012). These approaches allow us to estimate voxel-averaged microstructural information of the *in-vivo* brain that used to be accessible only via *ex-vivo* histology.

One of the promising contrasts for the new technology is magnetic susceptibility, measured by T_2_* decay and/or phase evolution (Duyn et al., 2007; Haacke et al., 2005). The contrast was utilized to reveal laminar structures in cortices (e.g., stria of Gennari (Duyn et al., 2007)), tiny nuclei in subcortices (e.g., nigrosome 1 and 4 (Sung et al., 2018) and subthalamic nuclei (Deistung et al., 2013)), and variations of iron concentration in superficial white matter (Kirilina et al., 2020).

In the brain, the susceptibility contrast has been demonstrated to originate primarily from two sources: iron and myelin (Duyn and Schenck, 2017). These substances have essential roles in normal brain functions. For example, iron is known to contribute to the synthesis of myelin, DNA, and neurotransmitters (Ward et al., 2014). Changes in iron homeostasis are related to the pathogenesis of multiple sclerosis (MS), Alzheimer’s diseases (AD), and Parkinson’s disease (Stephenson et al., 2014; Zecca et al., 2004). For myelin, production and degradation have shown to be involved in development (Sowell et al., 2003), neuro-plasticity (McKenzie et al., 2014), and neurodegenerative diseases such as MS and leukodystrophy (Compston and Coles, 2008; Nave, 2010). As a result, the two substances have been suggested as important biomarkers for neurological disorders.

Interestingly, iron and myelin have the opposite magnetic susceptibility characteristics: paramagnetic iron vs. diamagnetic myelin. This difference may provide an opportunity to differentiate the substances. However, measuring the individual concentration is still challenging because they co-exist in most of the brain regions and collectively determine the susceptibility contrasts (Schenck, 1996; Yablonskiy and Haacke, 1994). Efforts have been made to separate the contributions of iron and myelin. Schweser et al. tried to estimate iron and myelin concentrations from quantitative susceptibility mapping (QSM) and R_2_* using a linear equation model, which required an assumption of a magnetization transfer contrast reflecting myelin (Schweser et al., 2011b). Stüber et al. proposed to infer iron and myelin concentrations via a multivariate linear regression analysis between R_1_ and R_2_* vs. iron and myelin concentrations measured by proton-induced X-ray emission (Stüber et al., 2014). Similar approaches were applied to track the signatures of iron accumulation and demyelination in MS patients (Elkady et al., 2018, 2017; Pontillo et al., 2021). In these studies, however, the relation between the MRI measurements and the two substances were determined empirically rather than via biophysical modeling (see Discussion).

In this work, we formulate a new biophysical model that explains the individual contribution of paramagnetic (e.g., iron) and diamagnetic (e.g., myelin) susceptibility sources to MRI signals. By exploiting the model, we develop a new susceptibility source separation method, *χ*-separation, which estimates the individual concentration of the two sources even when they co-exist in a voxel. Our biophysical model and *χ*-separation method are validated using a numerical simulation and an experimental phantom. Then, the *χ*-separation method is applied to an *ex-vivo* human brain specimen to qualitatively compare the *χ*-separation results with an iron image from laser ablation-inductively coupled plasma-mass spectrometry (LA-ICP-MS) and a myelin image from Luxol fast blue (LFB) stain. Finally, the method is applied to the *in-vivo* human brain, revealing histologically well-known features of iron and myelin. In MS patients, the *χ*-separation results suggest to characterize lesion types, demonstrating the feasibility of applying the method for clinical research.

## 2. Methods

### 2.1. Biophysical model for χ-separation

When a susceptibility source is positioned in a magnetic field, the field within and around the source is perturbed according to Maxwell’s equations (Fig. 1). This field perturbation creates variations in the resonance frequency of MRI. The frequency shift (Δ*f*), originating from a bulk magnetic susceptibility distribution (*χ***(*r*)**), has been modeled as follows (Salomir et al., 2003):

**Figure 1.**
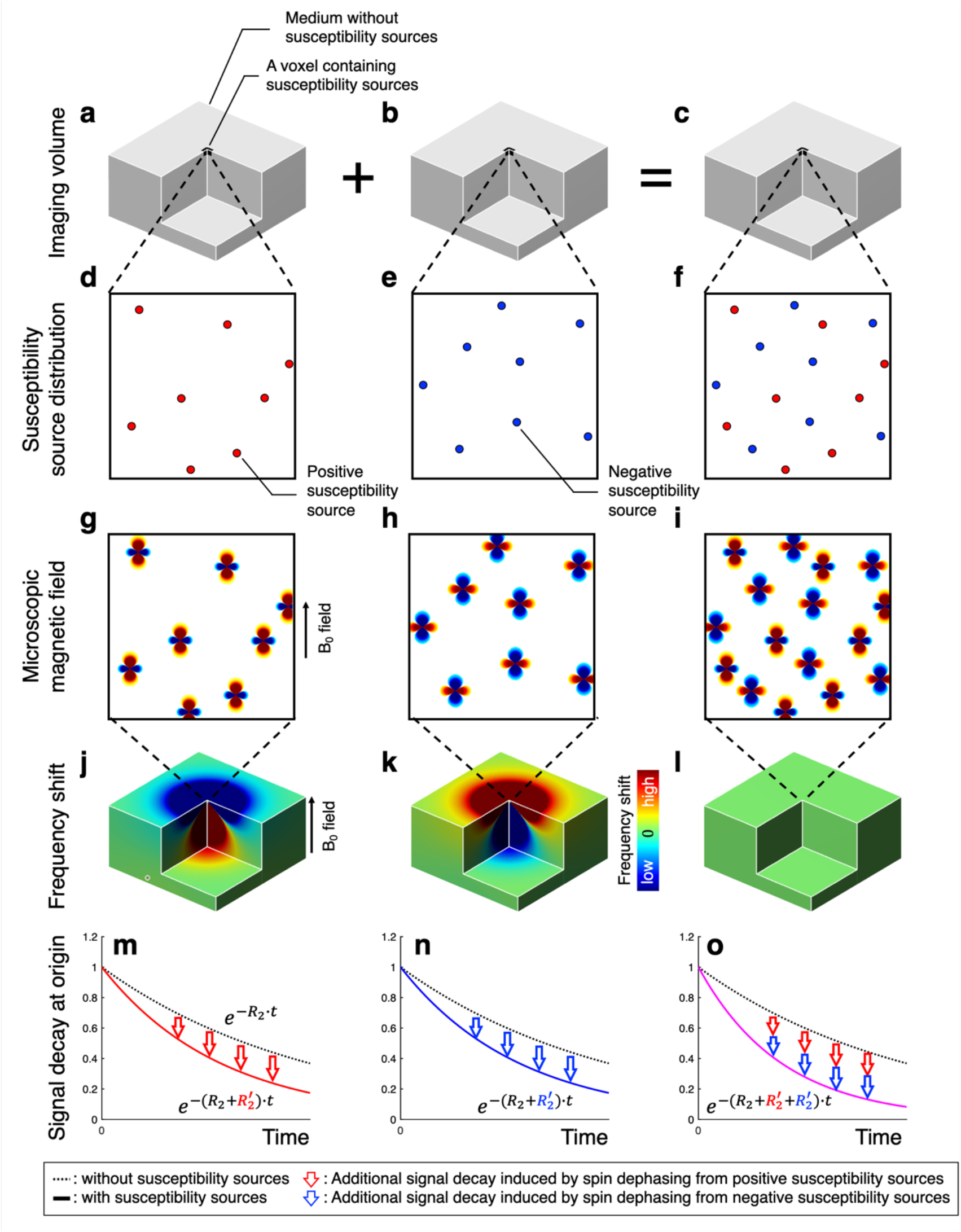
Conceptual illustration of MRI resonance frequency shift and transverse signal decay from positive and negative susceptibility sources. **a-c**, Imaging volume with a voxel at the origin containing randomly distributed spherical susceptibility sources. The sources have positive susceptibility in the first column, negative susceptibility in the second column, and both positive and negative susceptibility in the third column. **d-f**, Microscopic view of the susceptibility source distribution. **g-i**, Magnetic field perturbation induced by the susceptibility sources when the B_0_ field is applied. **j-l**, Voxel-averaged frequency shift in the imaging volume. The frequency shift is zero when the same amounts of the positive and negative susceptibility sources exist in the voxel (**l**). **m-o**, Transverse signal decay with irreversible (R_2_) and reversible (R_2_’) transverse relaxation rates in the voxel at the origin. The voxel containing both positive and negative susceptibility sources show R_2_’ as the sum of R_2_’ from the positive-source-only voxel and the negative-source-only voxel (**o**).

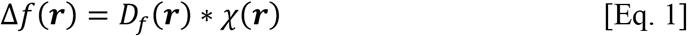

where ***r*** is the position vector of a voxel, *D*_*f*_ is a field perturbation kernel, and ∗ denotes a convolution operation. The kernel can be analytically derived from Maxwell’s magneto-static equations (Marques and Bowtell, 2005; Salomir et al., 2003) and has a well-known field pattern of a magnetic dipole. In conventional QSM, Δ*f* is obtained from the phase of an MRI signal, and then *χ* is estimated by the deconvolution of *D*_*f*_ from Δ*f* (Rochefort et al., 2008; Shmueli et al., 2009).

If a voxel contains both paramagnetic and diamagnetic susceptibility sources, the bulk magnetic susceptibility can be expressed as the sum of positive susceptibility (i.e., *χ*_*pos*_ > 0) and negative susceptibility (i.e., *χ*_*neg*_ < 0) of the voxel. Then, the frequency shift can be expressed as the signed sum of the effects of two susceptibility sources. Therefore, Eq. 1 can be written as follows:

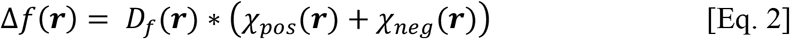

where *χ*_*pos*_ and *χ*_*neg*_ are the bulk susceptibility of the voxel from the positive and negative susceptibility sources, respectively.

The decay of a transverse MRI signal has been described by two different time constants, R_2_ and R_2_’, which are referred to as irreversible and reversible transverse relaxation rates, respectively. The R_2_’ relaxation originates primarily from magnetic susceptibility sources when ignoring chemical exchange and chemical shift effects. In the static dephasing regime, which postulates low diffusivity and low susceptibility source concentration, the decay of a voxel can be modeled as follows (Yablonskiy and Haacke, 1994) :

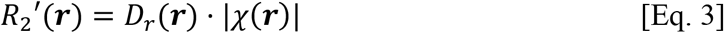

where *D*_*r*_ is a relaxometric constant between R_2_’ and susceptibility. This equation suggests that R_2_’ is linearly proportional to the concentration of susceptibility sources regardless of the sign of them. The relaxometric constant can be estimated by the ratio of R_2_’ to absolute susceptibility.

The R_2_’ model in Eq. 3 can be extended to consider a voxel that contains both positive and negative susceptibility sources (see Supplementary Note):

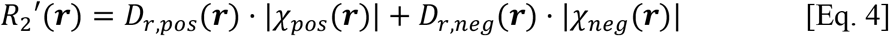

where *D*_*r,pos*_ and *D*_*r,neg*_ are the relaxometric constants for the positive and negative susceptibility sources, respectively. An important implication of this equation is that R_2_’ is determined by the (weighted) absolute sum of the effects of the two susceptibility sources.

Using Eqs. 2 and 4, the effects of magnetic susceptibility on a complex MRI signal can be modeled as follows:

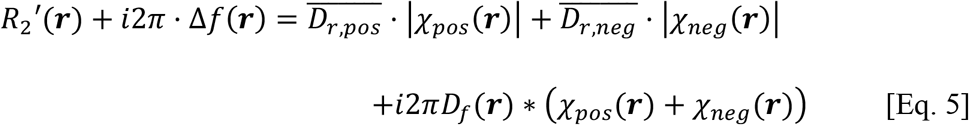

where 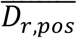 and 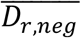 are nominal relaxometric constants for the positive and negative susceptibility sources, respectively, assuming spatial invariance. Eq. 5 was reformulated as a minimization problem and was solved iteratively using a conjugate gradient descent algorithm. For details of implementation, see Supplementary Information. Note that this model assumes susceptibility is the primary source for frequency shift and R_2_’, while ignoring susceptibility anisotropy (Lee et al., 2010) or water compartmentalization (Wharton and Bowtell, 2015, 2012).

### 2.2. Monte-Carlo simulation

To validate the two susceptibility models (Eqs. 2 and 4), and the *χ*-separation method (Eq. 5), a Monte-Carlo simulation was performed. A total of nine segments were designed to have the nine different combinations of susceptibility source compositions (positive-source-only, negative-source-only, and sum of the positive and negative sources) and susceptibility concentrations (±0.0125, ±0.025, and ±0.0375 ppm) (Fig. 2a-b). Each segment had 26 × 26 × 32 voxels, in which existed a cylinder (diameter: 8 voxels; length: 32 voxels) containing susceptibility sources (radius: 1 µm; susceptibility: +520 (Schenck, 1996) or −520 ppm). The positions of the susceptibility sources were randomly determined in the segment. The field perturbation from the susceptibility sources was calculated using Maxwell’s equation (Li and Leigh, 2004), assuming the B_0_ field of 3 T. The longitudinal axis of the cylinders was assigned to be perpendicular to the B_0_ field. In each voxel, 1000 protons performed random walks, following Gaussian diffusion with the diffusion coefficient of 1 µm^2^/ms. Each proton had a unit magnetization and R_2_ of 10 Hz. In each time step (100 µs), the phase of the proton magnetization was accumulated by the susceptibility-induced field. Then, the voxel-averaged signal was calculated for the echo time (TE) of 2.6 ms to 27.1 ms with the echo spacing of 4.9 ms. This simulation was repeated 444 times with randomly repositioned protons and susceptibility sources, and the results were averaged for each voxel. After that, the magnitude signal decay was fitted to an exponential model to estimate R_2_*, whereas the phase evolution was fitted to a linear model to measure frequency shift. The R_2_’ map was calculated by subtracting the R_2_ from the R_2_* map. Other details of the simulation are described in Supplementary Information. The simulation was performed using 12 GPUs (Nvidia GTX 1080Ti, Santa Clara, CA), and the total simulation time was 21 days.

**Figure 2.**
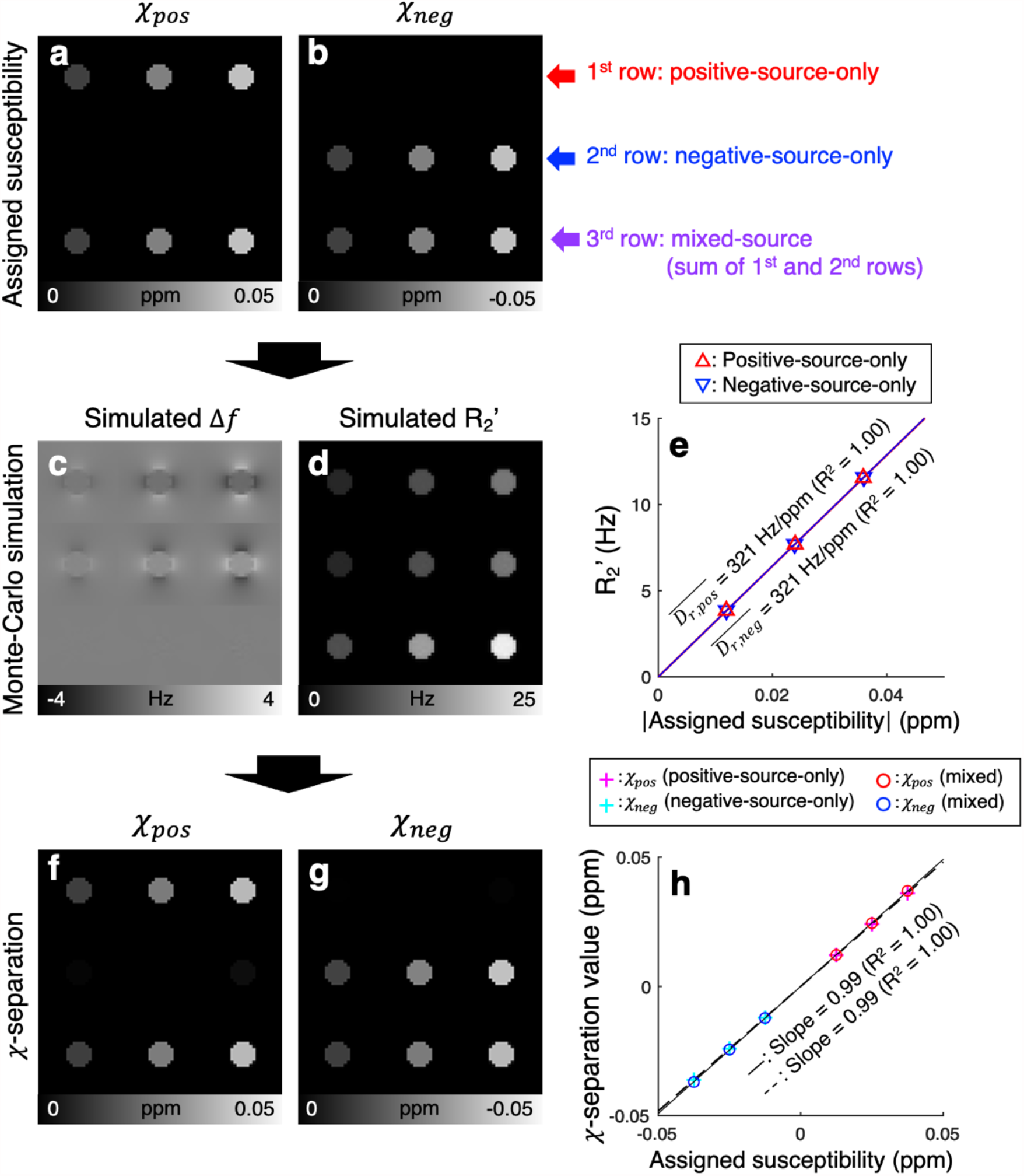
Validation of *χ*-separation using the Monte-Carlo simulation. **a-b**, Positive and negative susceptibility maps assigned for the simulation. **c-d**, Frequency shift (Δ*f*) and R_2_’ maps generated from the simulation. **e**, Absolute values of the assigned susceptibility vs. R_2_’ (red upper triangles: positive-source-only cylinders; blue lower triangles: negative-source-only cylinders). Both of the relaxometric constants are 321 Hz/ppm (R^2^ = 1.00). **f-g**, Positive and negative susceptibility maps reconstructed from *χ*-separation. The *χ*-separation results demonstrate a successful reconstruction of the assigned susceptibility. **h**, Assigned susceptibility vs. *χ*-separation results (magenta crosses: positive susceptibility in the positive-source-only cylinders; cyan crosses: negative susceptibility in the negative-source-only cylinders; red circles: positive susceptibility in the mixed-source cylinders; blue circles: negative susceptibility in the mixed-source cylinders). The susceptibility values from *χ*-separation match well with the assigned susceptibility values (regression results of assigned vs. single-source-only cylinder measurements: slope = 0.99 with R^2^ = 1.00, dashed line; regression results of assigned vs. mixed-source cylinder measurements: slope = 0.99 with R^2^ = 1.00, solid line).

The relaxometric constants were determined by the slope of the linear regression line of the averaged R_2_’ values with respect to the absolute values of the assigned susceptibility in the positive-source-only cylinders for 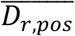 and the negative-source-only cylinders for 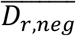. After that, the *χ*-separation algorithm was applied to each segment, reconstructing the positive and negative susceptibility maps. The mean susceptibility value of each cylinder was compared with the assigned value. For analysis, linear regression was performed for the susceptibility values from the positive-source-only and negative-source-only cylinders and for those from the mixed-source cylinders.

### 2.3. Phantom experiment

For the phantom experiment, a large cylindrical container (diameter: 120 mm, length: 92 mm) was filled with 1.5% agarose gel (Agarose ME, Daejung, Siheung, Republic of Korea). Before the gel hardened, a plastic cast was positioned to form a 3 × 3 hollow cylinder array (cylinder diameter: 12 mm, length: 46 mm). These hollow cylinders were filled with different compositions and concentrations of susceptibility sources mixed with 1.5% agarose gel. For the susceptibility sources, iron oxide (Fe_3_O_4_, Bangs Laboratories Inc., Fisher, IN, USA) was used for positive susceptibility, and calcium carbonate powder (CaCO_3_, Yakuri Pure Chemicals, Kyoto, Japan) for negative susceptibility. In the first-row cylinders of the array, the iron oxide-containing gel of 0.25, 0.50, and 0.75 mg/ml iron oxide concentrations was assigned. In the second row, the calcium carbonate powder-containing gel of 12.5, 25.0, and 37.5 mg/ml calcium carbonate concentrations was assigned. In the third-row cylinders, the mixtures of the iron oxide and calcium carbonate were created with the concentrations as the sum of the first two rows (Fig. 3c). To prevent precipitation of the susceptibility sources, an agarose solution was shaken before pouring into the hollow cylinder.

**Figure 3.**
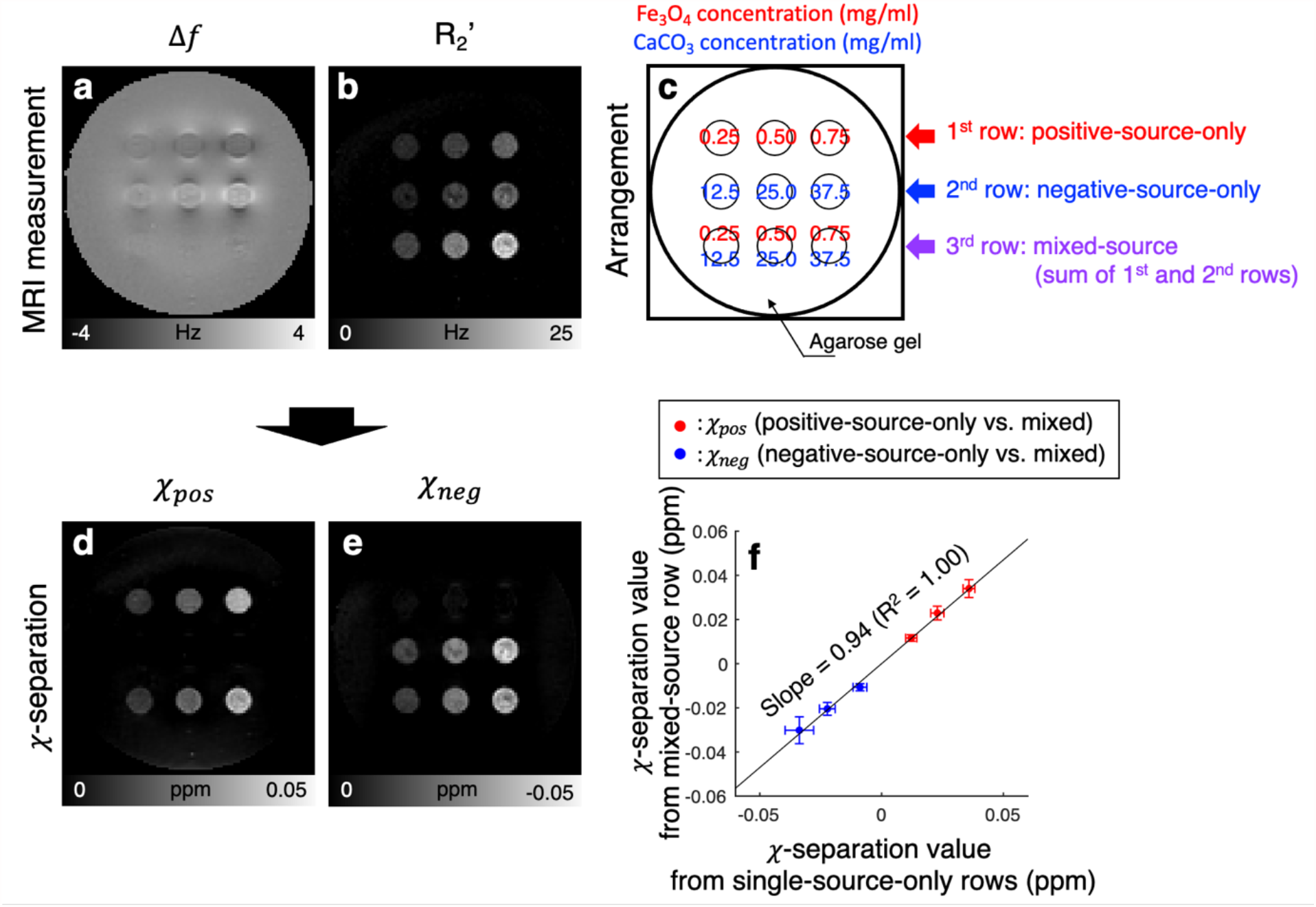
Validation of *χ*-separation using the phantom experiment. **a-b**, Frequency shift and R_2_’ maps from MRI measurements. **c**, Assigned susceptibility source concentrations (Fe_3_O_4_ for the positive susceptibility source and CaCO_3_ for the negative susceptibility source) **d-e**, Positive and negative susceptibility maps reconstructed from *χ*-separation. The results demonstrate successful separation of the two sources. **f**, Susceptibility measurements in the single-source-only cylinders vs. mixed-source cylinders (red dots: positive susceptibility in positive-source-only cylinders vs. mixed-source cylinders; blue dots: negative susceptibility in negative-source-only cylinders vs. mixed-source cylinders). The susceptibility values from the mixed-source cylinders match well with those from the single-source-only cylinders (regression results: slope = 0.94 with R^2^ of 1.00). Vertical and horizontal error bars in (**f**) indicate the standard deviation of the measurements in each cylinder.

The phantom was scanned at 3 T MRI (Siemens Tim Trio, Erlangen, Germany). The longitudinal axis of the small cylinders was placed perpendicular to the B_0_ field. For R_2_* and frequency shift, 3D multi-echo gradient-echo were acquired with the following parameters: field of view (FOV) = 192 × 192 × 120 mm^3^, voxel size = 1 × 1 × 2 mm^3^, repetition time (TR) = 60 ms, echo time (TE) = 2.6 ms, 7.5 ms, 12.4 ms, 17.3 ms, 22.2 ms, and 27.1 ms, bandwidth = 300 Hz/pixel, flip angle = 30°, and total acquisition time = 11.6 min. For R_2_, 2D multi-echo spin-echo data were acquired: FOV = 192 × 192 mm^2^, voxel size = 1 × 1 mm^2^, slice thickness = 2 mm, number of slices = 40, TR = 4000 ms, concatenation factor = 2, TE = 9 ms, 18 ms, 27 ms, 36 ms, 45 ms, 54 ms, 63 ms, 72 ms, 81 ms, and 90 ms, bandwidth = 300 Hz/pixel, and total acquisition time = 29.6 min.

After the acquisition, k-space data of all channels were reconstructed and then combined using the sum-of-squares for magnitude and MCPC-3D for phase (Robinson et al., 2011). To estimate the frequency shift, the multi-echo phase data were fitted to a linear model. Then, the result was spatially unwrapped (Cusack and Papadakis, 2002) and background-field-removed (Wu et al., 2012), producing a frequency shift map. From the frequency map, a conventional QSM map was reconstructed using the MEDI Toolbox (http://pre.weill.cornell.edu/mri/pages/qsm.html (Liu et al., 2018)). For R_2_* mapping, the magnitudes of multi-echo gradient echo signals were fitted to a mono-exponential decay function. For R_2_ mapping, the multi-echo spin-echo magnitude images were matched with a computer-simulated dictionary of spin-echo decay embedded with the stimulated echo correction (McPhee and Wilman, 2015). Then, an R_2_’ map was calculated by a voxel-wise subtraction of R_2_ from R_2_*. Negative R_2_’ values were refined to zero to enforce physics. The same reconstruction approach was performed for subsequent sections unless stated. The relaxometric constants were determined as the slope of the linear regression line of the mean R_2_’ with respect to the susceptibility values (estimated by the conventional QSM) in the positive-source-only cylinders for 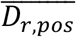 and the negative-source-only cylinders for 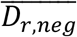 The voxels with extreme R_2_’ (> 25 Hz or < 2 Hz), which is most likely from the aggregation of the susceptibility sources, were excluded in the calculation. Finally, the positive and negative susceptibility maps were generated by *χ*-separation. To demonstrate successful susceptibility separation, the positive and negative susceptibility values in the mixed-source cylinders were compared to those in the single-source-only cylinders using a linear regression analysis.

### 2.4. Ex-vivo experiment

To demonstrate the specificity of *χ*-separation to iron and myelin in a biological tissue, an *ex-vivo* human brain specimen, containing the primary visual cortex (V1), was scanned. The results were compared with iron and myelin distributions measured by LA-ICP-MS and LFB myelin stain, respectively. The specimen was fixed in the 4% paraformaldehyde solution for four weeks after an autopsy, and then incubated in 0.1 M phosphate-buffered saline for seven days before being scanned at 7 T MRI (Siemens Terra, Erlangen, Germany). For the scan, 3D multi-echo gradient-echo data were acquired with the following parameters: FOV = 77 × 77 × 64 mm^3^, voxel size = 0.3 × 0.3 × 0.5 mm^3^, TR = 300 ms, TE = 3.9 ms, 10.4 ms, 16.9 ms, 23.5 ms, 30.0 ms, and 36.4 ms, flip angle = 45°, bandwidth = 300 Hz/pixel, two averages, and total acquisition time = 4.8 h. For 3D multi-echo spin-echo, the parameters had the same FOV and resolution as the gradient-echo acquisition, TR = 450 ms, TE = 14 ms, 28 ms, 42 ms, 56 ms, and 70 ms, bandwidth = 272 Hz/pixel, three averages, and total acquisition time = 10.6 h. The data were reconstructed in the same procedures as described in the phantom experiment. For *χ*-separation, the relaxometric constants from the health volunteers (see the next section) were used after scaling for the field strength difference (7 T vs. 3 T).

After the MRI scan, the specimen was paraffin-embedded and cut into 20-μm thick sections. The sections were utilized for the LFB myelin staining and LA-ICP-MS. For the myelin staining, the sections were deparaffinized and then immersed in a 0.1% LFB solution (LFB MBS 1.0 g (S3382, Sigma-Aldrich, St. Louis Luis, Mo, USA), 95% ethyl alcohol 1000 mL, and glacial acetic acid 5.0 mL (A6283, Sigma-Aldrich, St. Louis Luis, Mo, USA)) at 56 °C for 24 hours. The stain section was immersed in a 0.05% lithium carbonate solution (62470, Sigma-Aldrich, St. Louis Luis, Mo, USA) for tens of seconds for the differentiation (Fukunaga et al., 2010) and then imaged in a scanner with ×10 magnification (Axio Scan.Z1 slide scanner, Zeiss, Oberkochen, Germany). For the LA-ICP-MS, one of the sections was ablated using a 1030 nm femto-second pulsed laser system (J200 Femto iX, Applied Spectra Inc, West Sacramento, CA, USA) with the following parameters: laser spot diameter = 80 μm, line scan speed = 200 μm/s, repetition rate = 50 Hz, and raster spacing = 120 μm. The ablated tissue was transported by helium gas (0.9 L/min) with make-up argon gas (0.7 L/min) to a quadrupole ICP-MS device (iCAP TQ, ThermoFisher Scientific, Bremen, Germany), measuring ^56^Fe and ^13^C. The iron distribution (*Fe*_*LA*−*ICP*−*MS*_) was calculated as the signal intensity ratio of ^56^Fe to ^13^C to compensate for a system drift, assuming ^13^C as a reference (see Supplementary Fig. 2) (Lee et al., 2020). The chronological mass spectrum measurements were reshaped into a 2D image with the spatial resolution of 120 μm × 40 μm.

### 2.5. In-vivo experiment: healthy volunteers

Six healthy volunteers (age = 33.7 ± 14.6 years; 5 females and 1 male) were scanned at 3 T MRI (Siemens Tim Trio, Erlangen, Germany). All volunteers signed a written consent form approved by the institutional review board (IRB). The acquisition parameters for 3D multi-echo gradient-echo were FOV = 192 × 192 × 96 mm^3^, voxel size = 1 × 1 × 2 mm^3^, TR = 64 ms, TE = 2.9 ms, 7.5 ms, 12.1 ms, 16.7 ms, 21,3 ms, and 25.9 ms, bandwidth = 300 Hz/pixel, flip angle = 22°, parallel imaging factor = 2, and total acquisition time = 7.0 min. For 2D multi-echo spin-echo, the parameters were FOV = 192 × 192 mm^2^, voxel size = 1 × 1 mm^2^, slice thickness = 2 mm, number of slices = 40, TR = 3860 ms, TE = 15 ms, 30 ms, 45 ms, 60 ms, 75 ms, and 90 ms, bandwidth = 100 Hz/pixel, and total acquisition time = 12.4 min. The data were reconstructed as described in the phantom experiment. The relaxometric constant was estimated from five deep gray matter regions of interest (ROIs) that have primarily positive susceptibility: globus pallidus, putamen, caudate nucleus, substantia nigra, and red nucleus. The constant 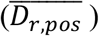 was determined by the slope of the linear regression line of the ROI-averaged R_2_’ values with respect to the susceptibility values (by the conventional QSM). For 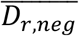 the same value as 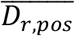 was used (see Discussion). After the reconstruction, the positive and negative susceptibility maps were compared with the histologically stained iron and myelin images from literatures (Drayer et al., 1986; Naidich et al., 2013; Schaltenbrand and Wahren, 1977).

### 2.6. In-vivo experiment: multiple sclerosis patients

Two MS patients (Patient 1: 34-year-old male; Patient 2: 23-year-old female; IRB-approved) were scanned using 3 T MRI (Philips Ingenia, Amsterdam, Netherlands). 3D multi-echo gradient-echo data were acquired with the following parameters: FOV = 216 × 216 × 144 mm^3^, voxel size = 0.5 × 0.5 × 2 mm^3^, TR = 42 ms, TE = 5.8 ms, 13.5 ms, 21.3 ms, 29.0 ms, and 36.7 ms, bandwidth = 255 Hz/pixel, flip angle = 17°, parallel imaging factor = 2.5 × 1.4, and total acquisition time = 5.1 min. For Patient 1, 2D multi-echo spin-echo data were acquired with the following parameters: FOV = 216 × 216 mm^2^, voxel size = 0.5 × 0.5 mm^2^, slice thickness = 2 mm, number of slices = 14, TR = 1800 ms, and TE = 10 ms, 20 ms, 30 ms, 40 ms, 50 ms, 60 ms, 70 ms, 80 ms, and 90 ms, bandwidth = 301 Hz/pixel, parallel imaging factor = 1.3, and total acquisition time = 7.8 min. For Patient 2, the same 2D multi-echo spin-echo data were acquired except for the following parameters: number of slices = 24, TR = 3300 ms, and TE = 20 to 100 ms with the echo spacing of 20 ms, bandwidth = 218 Hz/pixel, parallel imaging factor = 2.5, and total acquisition time = 7.2 min. For the clinical assessment of MS lesions, T_1_-weighted, T_2_-weighted, FLAIR, and gadolinium contrast-enhanced (CE) T_1_-weighted images were obtained (see Supplementary Information). The reconstruction was conducted in the same procedure as in the healthy volunteers except for the magnitude and phase generation which was performed by the scanner.

All data processing was performed using MATLAB (MATLAB 2020a, MathWorks Inc., Natick, MA, USA).

## 3. Results

### 3.1. Monte-Carlo simulation for validation

To demonstrate the validity of our susceptibility model (Eqs. 2 and 4) and *χ*-separation method, Monte-Carlo simulation is performed and the results are summarized in Fig. 2 and Supplementary Fig. 3. Nine cylinders with different combinations of susceptibility source compositions (first row: positive-source-only, second row: negative-source-only, and third row: sum of the positive and negative sources) and susceptibility concentrations are designed as shown in Fig. 2a-b. The resulting frequency shift maps demonstrate that the magnetic field perturbations around the positive-source-only cylinders and the negative-source-only cylinders show the dipole patterns as suggested in Eq. 1 (Fig. 2c). The R_2_’ maps, on the other hand, reveal localized R_2_’ within the source-containing voxels (Fig. 2d). The cylinders in the last row, containing the mixtures of the positive and negative susceptibility sources, demonstrate zero frequency shift in the frequency shift maps, while the R_2_’ maps report the contrasts proportional to the absolute sum of the positive and negative susceptibility (Supplementary Fig. 3). These observations validate our susceptibility model, which states that the signed sum of *χ*_*pos*_ and *χ*_*neg*_ determines the frequency shift while the absolute sum of them decides R_2_’. The positive and negative relaxometric constants are measured as 321 Hz/ppm (Fig. 2e). When *χ*-separation is applied to the Monte-Carlo simulated frequency shift and R_2_’ maps, the positive and negative susceptibility maps demonstrate successful separation of the two sources as demonstrated in Fig. 2f-g. The linear regression result confirms the accuracy of the *χ*-separation results (assigned susceptibility vs. measured susceptibility in the single-source-only cylinders: slope = 0.99, R^2^ = 1.00, dashed line in Fig. 2h; assigned susceptibility vs. measured susceptibility in the mixed-source cylinders: slope = 0.99, R^2^ = 1.00, solid line in Fig. 2h; slight underestimation from the numerical errors in the dipole kernel calculation).

### 3.2. Phantom experiment for validation

A phantom experiment using the positive (Fe_3_O_4_) and negative (CaCO_3_) susceptibility sources further consolidates the validity of our model and *χ*-separation. Fig. 3c shows the arrangement of the phantom with the different compositions of Fe_3_O_4_ and CaCO_3_ (first row: positive-source-only, second row: negative-source-only, and third row: sum of the positive and negative sources). The frequency shift map demonstrates the opposite field patterns between the positive (first-row) and negative (second-row) susceptibility sources (Fig. 3a). The third row, containing both sources, shows no conspicuous frequency shift, suggesting the signed sum of the susceptibility sources determines the frequency shift. On the other hand, R_2_’ reports dependence not on the signed sum but on the absolute sum of the susceptibility sources (Fig. 3b and Supplementary Fig. 4). The relaxometric constants are measured as 275 Hz/ppm (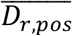; R^2^ = 1.00) for the positive source and 291 Hz/ppm (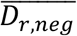; R^2^ = 0.83) for the negative source (Supplementary Fig. 5). When *χ*-separation is applied, the results demonstrate successful separation of the two sources, revealing the positive susceptibility in the first- and third-row cylinders and the negative susceptibility in the second- and third-row cylinders (Fig. 3d-e). The linear regression results in Fig. 3f show that the susceptibility values in the positive-source-only and negative-source-only cylinders agree with those in the mixed-source cylinders, confirming the validity of *χ*-separation (slope = 0.94; R^2^ = 1.00).

### 3.3. Ex-vivo brain specimen

When *χ*-separation is applied to an *ex-vivo* brain specimen containing V1, the positive and negative susceptibility maps reveal great similarities to the iron image from LA-ICP-MS and the myelin image from LFB myelin staining, respectively (Fig. 4; see Supplementary Fig. 6 for zoomed-in) despite complex microscopic environment of the brain (see Discussion). Overall, the cortex shows higher positive susceptibility and lower negative susceptibility than white matter (Fig. 4c-d). These patterns are in good agreement with the iron and myelin images in Fig. 4e and f. In V1, which is near the calcarine fissure outlined by the green lines, distinct laminar patterns are captured in the positive and negative susceptibility maps. The positive susceptibility map (Fig. 4c) shows three distinct layers, which reveal very low intensity in the superficial layer, an intensity peak in the middle layer (yellow arrowheads), and high intensity in the deep layer. These patterns are well-recognized in the iron image (Fig. 4e). In both images, the intensity peak disappears in the secondary visual cortex (V2). Compared to the positive susceptibility map, the negative susceptibility map (Fig. 4d) demonstrates a different pattern that illustrates low intensity in the superficial layer, an intensity peak in the middle layer, and low intensity in the deep layer, agreeing with the myelin histology image (Fig. 4f). The distinct peak in the middle layer of both susceptibility maps (Fig. 4c-d, yellow arrowheads) coincides with the previously reported histological characteristics of the stria of Gennari, featuring the co-existence of high concentrations of iron and myelin (see Supplementary Fig. 6 for a zoomed-in image) (Fukunaga et al., 2010). Intensity variations in the susceptibility maps are also observed in white matter and are in accordance with the iron and myelin images. In particular, stratum sagittale internum (purple squares in Fig. 4c-d) reports higher positive susceptibility and lower negative susceptibility than the surrounding white matter regions (red triangles: optic radiation; green stars: forceps), agreeing with the iron and myelin images (Fig. 4e-f) and previous reports (Duyn and Schenck, 2017; Sachs, 1893). On the other hand, these histological features are scantily observable or interpretable in the conventional contrasts (Fig. 4a-b and Supplementary Fig. 7).

**Figure 4.**
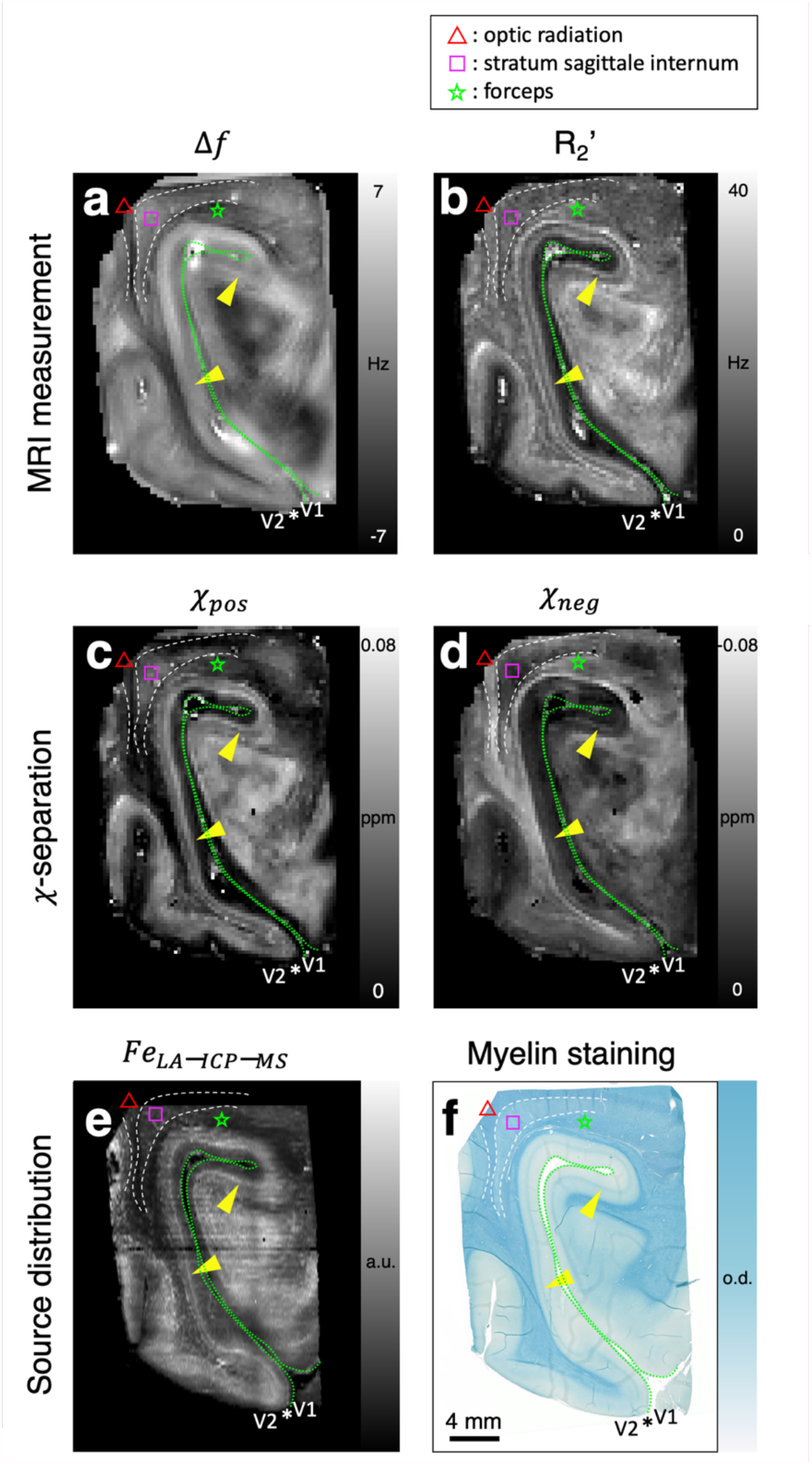
Comparison between *χ*-separation maps vs. iron and myelin images in the human brain specimen containing V1. **a-b**, Frequency shift and R_2_’ maps from MRI measurements. **c-d**, Positive (*χ*_pos_) and negative susceptibility (*χ*_neg_) maps from *χ*-separation. **e**, Iron image from LA-ICP-MS **(*Fe*_*LA*-*ICP*-*MS*_)**. **f**, Myelin image from LFB myelin staining. The positive susceptibility map reveals similar contrasts to the iron image whereas the negative susceptibility map shows similar features to the myelin image. Yellow arrowheads point to the stria of Gennari that shows high concentrations of iron and myelin (see Supplementary Fig. 6 for zoomed-in). Asterisks indicate V1-V2 boundaries. The pial surfaces of the calcarine fissure are delineated by green dashed lines. Red triangles, purple squares, and green stars mark optic radiation, stratum sagittale internum, and forceps, respectively. These white matter fibers show different concentrations of iron and myelin (**e-f**) which correspond well with the *χ*-separation results (**c-d**).

### 3.4. Healthy volunteers

When *χ*-separation is applied to healthy volunteers, the positive and negative susceptibility maps (Fig. 5 for a whole-brain slice and Fig. 6 for a few zoomed-in regions) render well-known characteristics of iron and myelin distributions that are not available in conventional contrasts (Fig. 5a-c, Fig. 5g-h, and Supplementary Fig. 8-9), suggesting the utility of *χ*-separation *in vivo*. For example, the deep gray matter regions show not only high positive susceptibility but also low negative susceptibility concentrations (yellow arrows in Fig. 5d-e), agreeing with high iron and low myelin concentrations in the areas (Drayer et al., 1986; Schaltenbrand and Wahren, 1977). In the negative susceptibility map, white matter reveals high signal intensity. In particular, optic radiation and forceps major (green and blue arrows in Fig. 5e), which are known to have high concentrations of myelin (Sachs, 1893), demonstrate high negative susceptibility concentrations. On the other hand, cortical gray matter shows low signal intensity in the negative susceptibility map, agreeing with our myelin histology (Fig. 4f). Note that high negative susceptibility values are observed around large vessels (e.g., internal cerebral vein) in spite of the paramagnetic susceptibility of deoxyhemoglobin (see Discussion). The total susceptibility map (Fig. 5f) demonstrates good similarities to the conventional QSM map (Fig. 5c). When analyzed for a few ROIs, the results from the positive susceptibility map better explains the literature values of iron concentrations than those of the conventional QSM (Supplementary Fig. 10). The relaxometric constant is estimated as 137 Hz/ppm with R^2^ of 0.70 (Supplementary Fig. 11).

**Figure 5.**
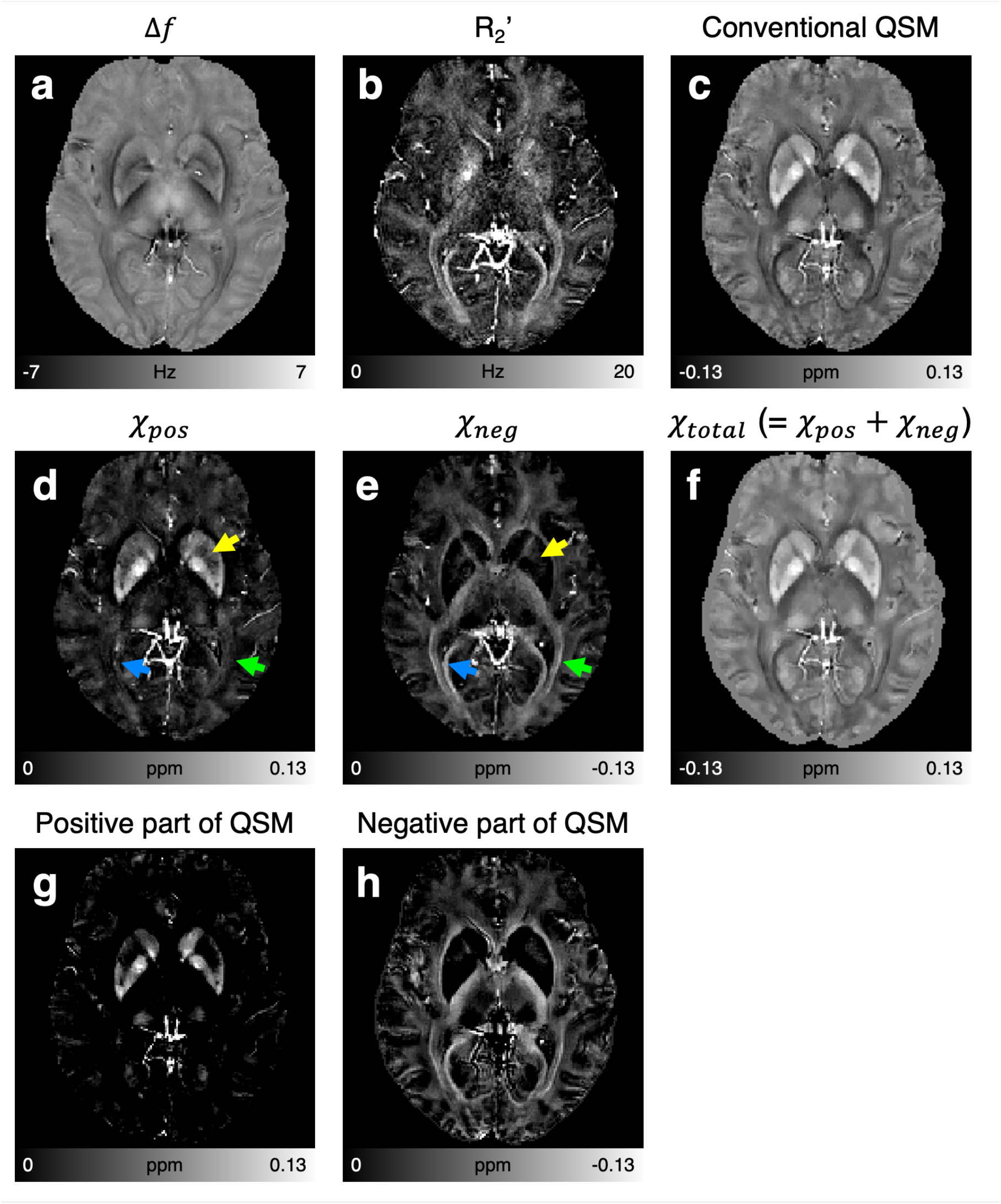
*In-vivo* results of a healthy volunteer. **a-c**, Frequency shift, R_2_’, and conventional QSM maps. **d-e**, Positive and negative susceptibility maps from *χ*-separation, reporting well-known iron and myelin distributions (e.g., high iron but low myelin concentrations in deep gray matter; high myelin concentrations in optic radiation and forceps major). Yellow, green, and blue arrows indicate deep gray matter, optic radiation, and forceps major, respectively. **f**, Total susceptibility map calculated as the sum of the positive and negative susceptibility maps. **g-h**, Positive and negative part maps of the conventional QSM map.

When the *in-vivo χ*-separation maps of the basal ganglia, thalamus, and midbrain areas are zoomed-in and compared with the histology images from the literatures (Drayer et al., 1986; Naidich et al., 2013; Schaltenbrand and Wahren, 1977), the positive and negative susceptibility maps match well with the iron and myelin histology results, respectively (Fig. 6). In basal ganglia (Fig. 6a-d), hyperintense positive susceptibility and hypointense negative susceptibility are observed in iron-rich but myelin-deficient regions including caudate nucleus (CN), globus pallidus (GP), and putamen (Put). On the other hand, hyperintensity in the negative susceptibility map (Fig. 6c) is observed in myelin-rich regions, depicting anterior limb of internal capsule (ALIC), external capsule (EC), fornix (Fx), and posterior limb of internal capsule (PLIC). In thalamus (Fig. 6e-h), the two sub-thalamic nuclei, pulvinar (Pul) and nucleus dorsomedialis (ND) where myelin concentrations are low (arrows in Fig. 6h), are identified in the negative susceptibility map (arrows in Fig. 6g). In particular, Pul shows high iron concentration (Fig. 6e), which is also observed in the positive susceptibility map (Fig. 6f). The rest of the thalamus reveals both positive and negative susceptibility sources that are supported by the iron and myelin histology images. In midbrain (Fig. 6i-l), red nucleus (RN) and substantia nigra (SN) show high iron but low myelin concentrations in histology. These variations are well-demonstrated in the positive and negative susceptibility maps (Fig. 6j-k) while the opposite characteristics being observed in crus cerebri (CrC).

**Figure 6.**
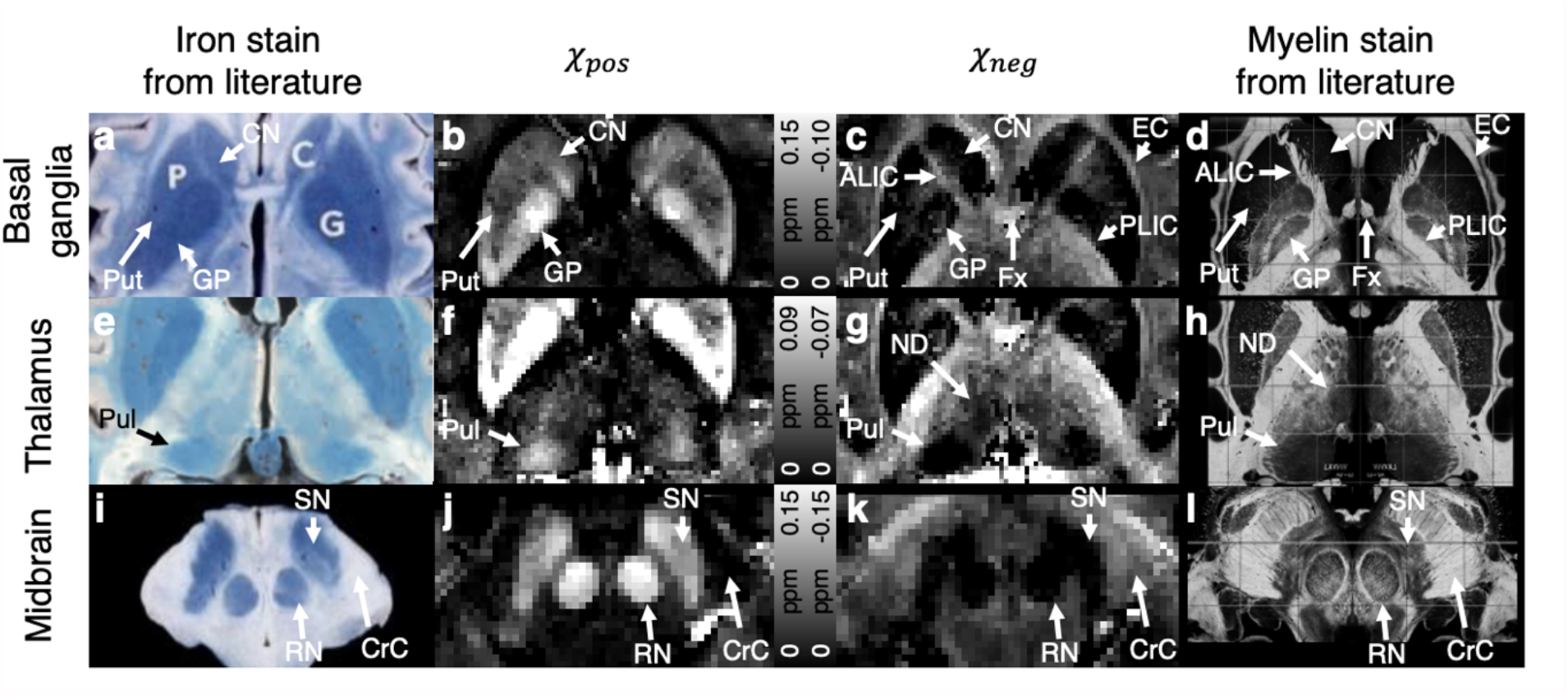
Comparison between *in-vivo χ*-separation results vs. iron and myelin histology from literatures. Zoom-in images of **(a-d)** basal ganglia, **(e-h)** thalamus, and **(i-l)** midbrain with **(a, e, i)** iron staining images, **(b, f, j)** positive susceptibility maps of *χ*-separation, **(c, g, k)** negative susceptibility maps of *χ*-separation, and **(d, h, l)** myelin staining images. Similar distributions are observed between the positive susceptibility maps and the iron staining images as well as the negative susceptibility maps and the myelin staining images. The *χ*-separation results reveal exquisite details of the brain anatomy that match with the iron and myelin histology: ALIC - anterior limb of internal capsule, CN - caudate nucleus, CrC - crus cerebri, EC - external capsule, Fx - fornix, GP - globus pallidus, ND - nucleus dorsomedialis, PLIC - Posterior limb of internal capsule, Pul - pulvinar, Put - putamen, RN - red nucleus, and SN - substantia nigra. Histological images are reproduced from Drayer et al. (Drayer et al., 1986) **(a, i)**, Naidich et al. (Naidich et al., 2013) **(e)**, and Schaltenbrand-Wahren (Schaltenbrand and Wahren, 1977) **(d, h, l)**. [Reprinted/Modified/Adapted from Burton Drayer et al, MRI of Brain Iron, American Journal of Roentgenology, 147:103-110, Copyright©1986, copyright owner as specified in the American Journal of Roentgenology]. For visualization, the myelin staining images were intensity-inverted and replicated with the left-right symmetry.

### 3.5. Multiple sclerosis patients

The results from two MS patients in Fig. 7 demonstrate the application of *χ*-separation in characterizing MS lesions. A lesion in Patient 1 (Fig. 7a, white box) renders rim-shaped hyperintensity in the positive susceptibility map (Fig. 7b) and overall hypointensity in the negative susceptibility map (Fig. 7c). These observations agree with the characteristics of the well-known “iron-rim lesion,” which has iron accumulation at the lesion boundary and demyelination in the lesion core (Bagnato et al., 2011). Different from this lesion, a lesion in Patient 2 (Fig. 7g, white box) show hypointensity in both positive and negative susceptibility maps (Fig. 7h-i). These patterns are in line with the histology of another popular lesion subtype, which has demyelination and iron depletion in the lesion (Hametner et al., 2013). In the conventional contrasts (Fig. 7d-f and j-l and Supplementary Fig. 12), however, these interpretations are not possible because the contrasts measure the combined effects of iron and myelin. A few artifacts including negative susceptibility around large veins, R_2_’ errors in the frontal lobe influencing both positive and negative susceptibility maps, and well-known streaking artifacts of QSM producing susceptibility variations near the iron-rim lesion of Patient 1 are noticed (see Discussion and Supplementary Fig. 14).

**Figure 7.**
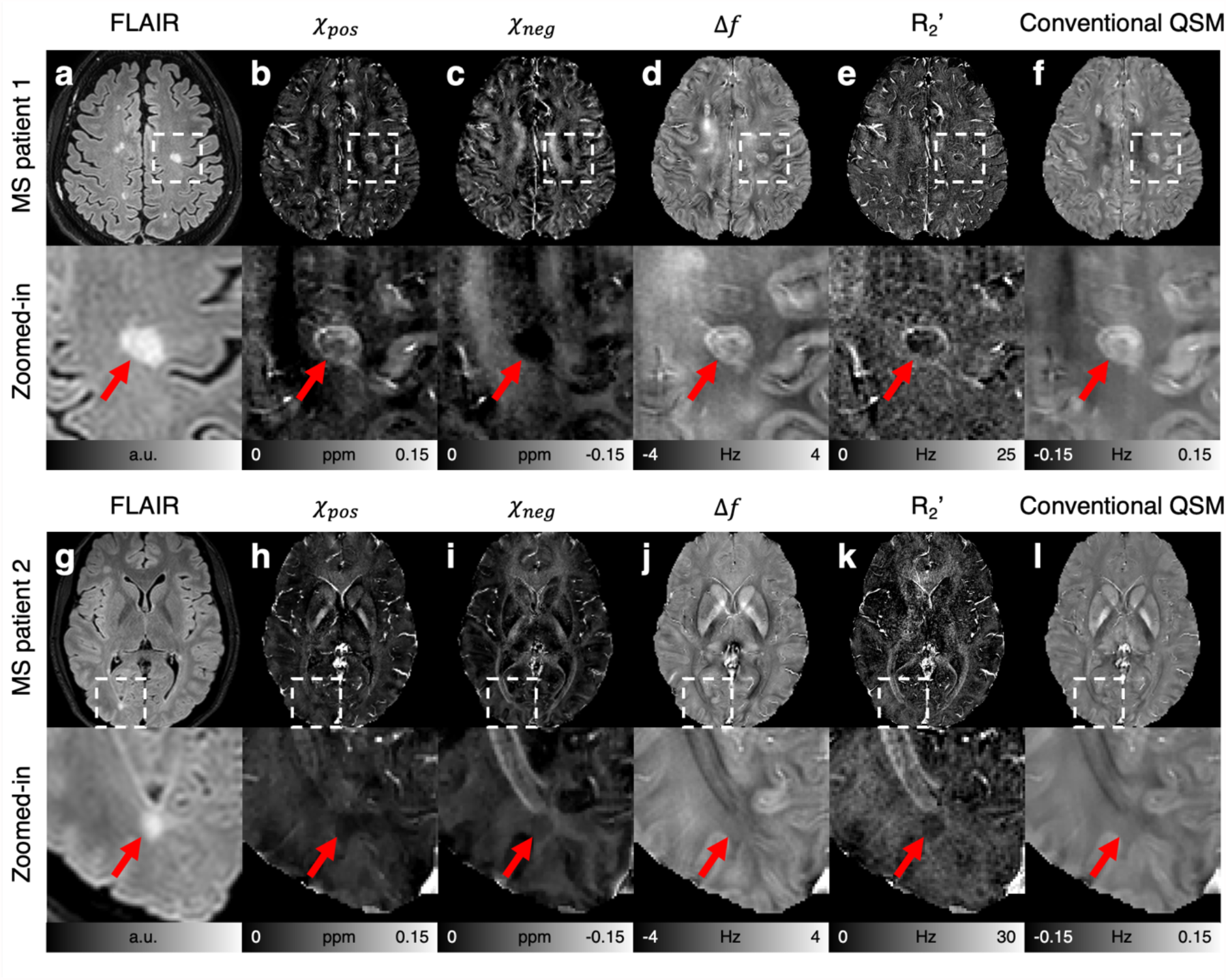
*In-vivo* results of two MS patients with characteristic lesions. **a and g**, FLAIR images. **b and h**, Positive susceptibility maps. **c and i**, Negative susceptibility maps. **d and j**, Frequency shift maps. **e and k**, R_2_’ maps. **f and l**, Conventional QSM maps. The zoom-in images show the areas (white boxes) containing typical MS lesions (MS patient 1: demyelinated iron-rim lesion; MS patient 2: demyelinated lesion with iron depletion). Red arrows indicate the location of the lesions in the FLAIR images.

## 4. Discussion

The newly proposed method, *χ*-separation, reveals the details of brain anatomy that are not easily visible in conventional MR images. In particular, the clear delineation of subthalamic nuclei and midbrain structures (Fig. 6) may provide valuable information in such applications as deep brain stimulation (Papavassiliou et al., 2004) and focused ultrasound surgery (Lipsman et al., 2013). Similarly, *χ*-separation images can be used as inputs for multi-contrast brain parcellation (Glasser et al., 2016), providing a subject-specific localization of subthalamic nuclei and midbrain structures. The localization may be used to detect the volume change of subthalamic nuclei in psychosis (Alemán-Gómez et al., 2020). In MS research, the iron and myelin information may improve the understanding of the disease pathogenesis and lesion characterization, which are limited by the co-variation of iron and myelin in lesions and surrounding areas (Wiggermann et al., 2017; Wisnieff et al., 2015). Another application is AD where amyloid plaque, which is diamagnetic (Gong et al., 2019), is one of the histopathological hallmarks of the disease. The magnetic property of the plaque is changed to be paramagnetic when iron accumulates in the plaque (Telling et al., 2017). The resulting paramagnetic property of the amyloid plaque has been captured in R_2_* and QSM (Jack et al., 2004; Tuzzi et al., 2020). The *χ*-separation method may distinguish paramagnetic iron from diamagnetic amyloid plaque, potentially introducing a new AD imaging biomarker.

In a few previous studies, methods were proposed to estimate iron and myelin concentrations from MRI measurements (Schweser et al., 2011a, 2011b; Stüber et al., 2014). These studies, however, developed empirically determined linear relationships between MRI measurements vs. iron and myelin, and/or utilized another MRI contrast to substitute myelin measurement. In the work by Schweser et al., regional iron was quantified by removing myelin contribution from QSM. In their work, myelin was estimated by a magnetization transfer contrast (Schweser et al., 2011a), which has shown to have a moderate correlation with myelin (Weijden et al., 2021). In their subsequent conference abstract, they tried to estimate iron and myelin distributions from QSM and R_2_* by suggesting another model in which R_2_* and QSM were assumed to be a linear function of iron concentration and the magnetization transfer contrast (Schweser et al., 2011b). Stüber et al. proposed a new approach that inferred iron and myelin concentrations from R_2_* and R_1_ via a linear equation between the two quantities vs. iron and myelin concentrations measured by proton-induced X-ray emission (Stüber et al., 2014). These studies are based on empirical relationships and may have limited accuracy, potentially hampering wide applications of the methods. On the other hand, our *χ*-separation model is built upon a biophysical model and has successfully separated the sources not only in the brain areas with a single dominant source (e.g., iron-rich but myelin-deficient globus pallidus) but also in the areas with both sources co-existing (e.g., stria of Gennari and subregions in thalamus).

In this work, the positive and negative susceptibility maps are regarded as markers for iron and myelin, respectively, assuming the two are the only sources of magnetic susceptibility in the brain. The assumption is supported by previous studies (Duyn and Schenck, 2017; Hametner et al., 2018). Although rare, however, additional susceptibility sources such as calcium (Chen et al., 2014) and copper (Fritzsch et al., 2014) may exist particularly in pathological conditions. Therefore, interpreting the results requires caution.

White matter has a complex microstructural geometry, which is not captured in our model, potentially creating errors in the susceptibility estimation. Despite this microstructural environment of the brain, our *ex-vivo* and *in-vivo* results show the potential of *χ*-separation in identifying the distribution of paramagnetic (i.e., iron) and diamagnetic (i.e., myelin) sources (Figs. 4 and 6). Recently, efforts have been made to build a susceptibility model of white matter, suggesting the susceptibility source (i.e., myelin) in the hollow cylinders (Sati et al., 2013; Wharton and Bowtell, 2012) with anisotropic effects and TE-dependent frequency shift (Lee et al., 2010; Wharton and Bowtell, 2015). Our model may be extended to incorporate this model and improve the accuracy when additional information such as fiber orientation is included. In such a model, the negative relaxometric constant can be formulated as a function of the fiber orientation.

Our model is established under the hypothesis that R_2_’ and frequency shift are determined by susceptibility concentration in the static dephasing regime. However, contrast sources such as chemical exchange may also contribute to frequency shift and/or R_2_’, although the effect size is smaller than that of susceptibility (Eun et al., 2020; Shmueli et al., 2011). Additionally, the validity of the static dephasing condition is influenced by water diffusivity, and the size, geometry, and characteristic frequency of susceptibility sources (Brammerloh et al., 2021; Duyn and Schenck, 2017) and may not strictly hold for brain regions with high iron concentrations (e.g., GP, RN, and SN in Fig. 6; (Yablonskiy et al., 2021). For example, if disease conditions alter diffusivity (Sugahara et al., 1999) or sequence parameters influence diffusion effects (e.g., echo space in spin echo (Muller et al., 1991; Ye and Allen, 1995)), R_2_ and R_2_* measurements can be changed, affecting *χ*-separation results.

The relaxometric constants in the simulation and phantom experiment were estimated by changing the source concentrations and measuring the corresponding R_2_’ changes. On the other hand, the *in-vivo* constant was estimated from deep gray matter ROIs and was used for both positive and negative relaxometric constants due to the absence of knowledge on the source concentrations. Although assumed to be constant in our experiment, the *in-vivo* relaxometric constant may vary across brain regions, influenced by the size, geometry, and characteristic frequency of susceptibility sources (Gossuin et al., 2004; Schenck and Zimmerman, 2004; Taege et al., 2019). A further technique needs to be developed for an accurate measurement of the value. When we performed the *χ*-separation reconstruction of the computer simulation and *in-vivo* data for a range of the relaxometric constant (75% to 125% of the original value), the susceptibility estimations showed qualitatively similar contrasts (Supplementary Fig. 13), suggesting that our *in-vivo* results are at least qualitatively relevant.

In the Monte-Carlo simulation, the susceptibility measurements from *χ*-separation revealed slight underestimation when compared to the assigned susceptibility values (slopes of the linear lines: 0.99 in Fig. 2h). This underestimation can be explained by the numerical errors from a finite sampling of the dipole pattern, as demonstrated in a previous study (Zhou et al., 2017).

In the phantom experiment, the susceptibility sources, especially CaCO_3_, were not fully resolved in water, resulting in the aggregation of the sources. This aggregation is observed as a few spotty variations in the cylinders (Fig. 3) and maybe a source for the estimation errors in the *χ*-separation results (Fig. 3f; Supplementary Fig. 4d and e; Supplementary Fig. 5).

In Figs. 5 and 7, negative susceptibility values are observed in voxels near large vessels despite paramagnetic deoxyhemoglobin in the vessels. This error is due to the macroscopic field inhomogeneity effects from the vessels, creating non-local R_2_’. This condition violates our assumption of fully localized R_2_’ in Eq. 4, creating artifacts in the results. Similarly, areas with large B_0_ field inhomogeneity (e.g., frontal lobe) from imperfect shim reveal errors in the positive and negative susceptibility maps (Fig. 7; Supplementary Fig. 14). In addition, artifacts from inaccurate background field removal, dipole inversion (i.e., streaking artifacts), and fiber orientation effects, that are observed in conventional QSM, are also noticeable in the *χ*-separation results (Supplementary Fig. 14). Partly because of these issues and partly because of the challenges in spatial registration and quantitative evaluation of the histological measurements, the *χ*-separation results of the brain are qualitatively assessed. Further technical development is expected to reduce these artifacts and to improve the image quality of *χ*-separation.

In the *in-vivo* experiments, R_2_ was measured using a 2D sequence to avoid a long scan time whereas R_2_* was acquired in 3D, introducing discrepancy in the slice profiles between the two datasets. This mismatch may generate errors in *χ*-separation results, particularly in the regions of large field inhomogeneity. Further development of data acquisition methods such as efficient 3D R_2_ mapping (Nguyen et al., 2012) will be useful.

## 5. Conclusions

We have demonstrated that a new biophysical model, describing the individual contribution of the positive and negative susceptibility sources to MRI signals, enables us to generate the maps of positive and negative susceptibility sources. This new technology was validated using computer simulation and experimental phantom and applied to *ex-vivo* and *in-vivo* human brains, producing high quality images of iron and myelin non-invasively. The results of healthy volunteers illustrate exquisite details of histologically well-known information of iron and myelin distributions, localizing structures such as nucleus dorsomedialis and pulvinar in thalamus. The images from multiple sclerosis patients reveal distinct characteristics of a few multiple sclerosis lesions such as iron-rim lesion and demyelinated lesion with iron depletion. The acquisition of *χ*-separation data takes less than 20 min of MRI scan time, which can be further reduced with currently available fast imaging approaches, and, therefore, can be used as a practical tool for imaging microstructural information of the brain. Clinically, the technique could lead to *in-vivo* monitoring of pathogenesis of neurological diseases that entail changes in iron and myelin. It could also be applicable to neuroscience studies, exploring personalized localization of deep gray matter nuclei or identifying activity-induced myelin changes from neuroplasticity.

## Acknowledgments

This research was supported by the National Research Foundation of Korea (NRF-2021R1A2B5B03002783 and NRF-2017H1A2A1042711), INMC and IOER at Seoul National University, and Grant-in-Aid for Scientific Research (KAKENHI-19K22985) from Japan Society for the Promotion of Science. We thank Professor Norihiro Sadato for his support of this project and Institute for Basic Science Center for Neuroscience Imaging Research (IBS-R015-D1) for providing MRI time and professional technical support.

## Data and code availability

All data needed to evaluate the conclusions in the paper are present in the paper and/or the Supplementary Materials. Additional data related to this paper can be obtained from the authors upon request. The reconstruction codes and details of data acquisition related to this work will be available at https://github.com/SNU-LIST.

