## Supplementary Information for "*χ*-separation: Magnetic susceptibility source separation toward iron and myelin mapping in the brain"

### Table of contents

**Supplementary methods**

• Algorithm for $\chi$-separation

• Implementation details of Monte-Carlo simulation

• Imaging parameters for MS lesion assessment

- Supplementary Figure 1

**Supplementary figures**

• Supplementary Figures 2-14

**Supplementary note**

• Proof of $R_2'$ model in Eq. 4 for multiple susceptibility sources

**References**

#### Supplementary methods

##### *Algorithm for $\chi$-separation*

For the implementation of $\chi$-separation, Eq. 5 is formulated as the following minimization problem:

$$argmin_{\chi_{pos},\chi_{neg}} \left\|W_r \cdot \left\{R_2{}' - \left(\overline{D_{r,pos}} \cdot |\chi_{pos}| + \overline{D_{r,neg}} \cdot |\chi_{neg}|\right)\right\} + i2\pi \cdot W_f \cdot \left\{f - D_f * \left(\chi_{pos} + \chi_{neg}\right)\right\}\right\|_2^2 + reg(\chi_{pos},\chi_{neg})$$

$$subject\ to \quad \chi_{pos} \geq 0, \chi_{neg} \leq 0$$

where $W_f$ is a weight accounting for the signal-to-noise ratio of the gradient echo signals (Liu et al., 2011), and $W_r$ is a weight reducing the effects of unreliable $R_2'$:

$$W_r(\boldsymbol{r}) = \begin{cases} W_f(\boldsymbol{r})/10 & where\ R_2{}'(\boldsymbol{r}) > 30\text{ Hz}\ or\ R_2{}'(\boldsymbol{r}) < 1\text{ Hz}, \\ W_f(\boldsymbol{r}) & otherwise. \end{cases}$$

For the phantom data, the condition $R_2{}'(\boldsymbol{r}) < 1$ Hz was not applied in order to acquire signals from the agarose gel. The regularization term, $reg(\chi_{pos},\chi_{neg})$, is designed to reduce streaking artifacts in the results as suggested in the conventional QSM method (Liu et al., 2018).

$$reg(\chi_{pos},\chi_{neg}) = 2 \cdot \lambda_1 \left\|M_{Mag}(\nabla\chi_{total})\right\|_1 + \lambda_1 \left\|M_{R_2'}(\nabla\chi_{pos})\right\|_1 + \lambda_1 \left\|M_{R_2'}(\nabla\chi_{neg})\right\|_1 + \lambda_2 \left\|M_{CSF}(\chi_{pos} - \overline{\chi_{pos,CSF}})\right\|_2^2 + \lambda_2 \left\|M_{CSF}(\chi_{neg} - \overline{\chi_{neg,CSF}})\right\|_2^2$$

where $\lambda_1$ and $\lambda_2$ are regularization parameters, $\nabla$ is a gradient operation, $\chi_{total}$ is a total susceptibility map calculated as the sum of $\chi_{pos}$ and $\chi_{neg}$, $M_{Mag}$ is a binary edge mask from magnitude (Liu et al., 2011), $M_{R_2'}$ is a binary edge mask from $R_2'$, $M_{CSF}$ is a binary mask of ventricular cerebrospinal fluid (Liu et al., 2018), and $\overline{\chi_{\cdot,CSF}}$ is the mean positive (or negative) susceptibility in $M_{CSF}$ (Liu et al., 2018). All the other terms are the same as those in Eq. 5.

The minimization problem is solved iteratively using a conjugate gradient algorithm by simultaneously updating $\chi_{pos}$ and $\chi_{neg}$ in each iteration. If a susceptibility value violates the physical constraints (i.e., $\chi_{pos} \geq 0$ and $\chi_{neg} \leq 0$) during the iteration, it is forced to zero.

The iteration stops when the residual, which is defined as $\|\chi_{n+1} - \chi_n\|_2 / \|\chi_n\|_2$ where $\chi_n$ is the sum of $\chi_{pos}$ and $\chi_{neg}$ at the n$^{th}$ iteration, is less than 0.01, or when it reaches 30 iterations. The $\chi_{pos}$ and $\chi_{neg}$ maps are initialized as the solution of the following two linear equations:

$$D_{r,pos}(\boldsymbol{r}) \cdot |\chi_{pos}(\boldsymbol{r})| + D_{r,neg}(\boldsymbol{r}) \cdot |\chi_{neg}(\boldsymbol{r})| = R_2{}'(\boldsymbol{r}),$$

$$\chi_{pos} + \chi_{neg} = \chi_{conventional\ QSM}$$

where $\chi_{conventional\ QSM}$ is the reconstruction results of the conventional QSM (Liu et al., 2018).

*Implementation details of Monte-Carlo simulation*

To reduce the computational cost of the Monte-Carlo simulation, the number of the susceptibility sources was fixed to 10, while changing the voxel size of a segment to satisfy the susceptibility concentration. For example, a voxel size of 120 × 120 × 120 $\mu m^3$ was used to achieve the susceptibility concentration of 0.0125 ppm while 83 × 83 × 83 $\mu m^3$ was used for 0.0375 ppm. For the same reason, the protons were located only in the center slice. Then, the simulation results were appended along the z-direction, generating a 3D matrix (i.e., 26 × 26 × 32). This matrix was used for the susceptibility reconstruction.

The convention of a magnetic dipole in Eq. 2 assumes no frequency shift at the origin based on the Lorentz sphere (i.e., $D_f(\boldsymbol{r} = \boldsymbol{0}) = 0$) (Wang and Liu, 2015). However, studies have reported that the non-zero frequency shift exists at the origin (Ruh et al., 2018; Yablonskiy and Haacke, 1994). This frequency shift was compensated for the simulation results (Yablonskiy and Haacke, 1994).

*Imaging parameters for MS lesion assessment*

For the clinical assessment of MS lesions, the following four sequences were acquired in the MS patients.

- FLAIR image: FOV = 256 × 256 × 180 $mm^3$, voxel size = 1 × 1 × 1 $mm^3$, TR = 4800 ms, TE = 310 ms, bandwidth = 957 Hz/pixel, parallel imaging factor = 3 × 2, inversion time (TI) = 1650 ms, turbo factor = 167, refocusing angle = 30°, two averages, and total acquisition time = 5.6 min.
- $T_1$-weighted image: FOV = 256 × 256 × 176 $mm^3$, voxel size = 1 × 1 × 1 $mm^3$, TR = 5851 ms, TE = 1599 ms, bandwidth = 191 Hz/pixel, flip angle = 7°, parallel imaging factor = 2, TI = 1100 ms, turbo factor = 211, and total acquisition time = 5.4 min.
- $T_2$-weighted image: FOV = 240 × 240 × 188 $mm^3$, voxel size = 1 × 1 × 1 $mm^3$, TR = 2500 ms, TE = 300 ms, bandwidth = 609 Hz/pixel, parallel imaging factor = 2, turbo factor = 135, refocusing angle = 35°, and total acquisition time = 5.5 min.
- CE $T_1$-weighted image: FOV = 240 × 240 × 180 $mm^3$, voxel size = 1 × 1 × 1 $mm^3$, TR = 500 ms, TE = 30 ms, bandwidth = 755 Hz/pixel, flip angle = 80°, TI = 1650 ms, turbo factor = 30, refocusing angle = 35°, two averages, and total acquisition time = 3.9 min.

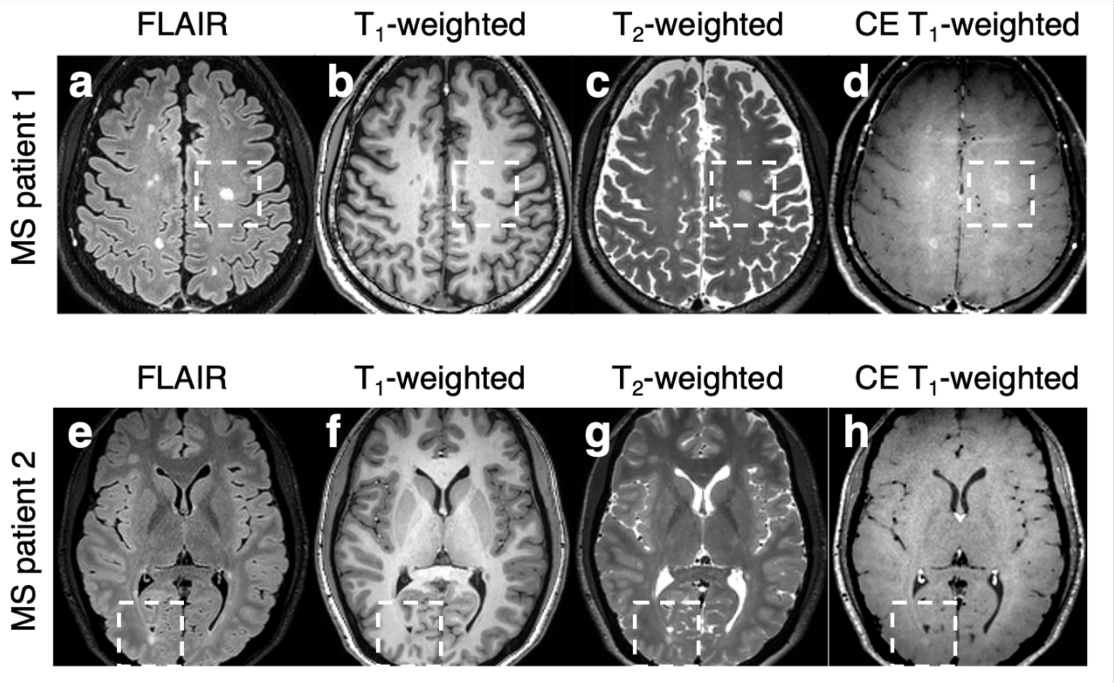

**Supplementary Figure 1. Images of the two MS patients (a-d from Patient 1; e-h from Patient 2) for the clinical assessment of MS lesions. a and e**, FLAIR images. **b and f**, $T_1$-weighted images. **c and g**, $T_2$-weighted images. **d and h**, CE $T_1$-weighted images. White boxes denote the areas of the MS lesions shown in Fig. 7.

**Supplementary figures**

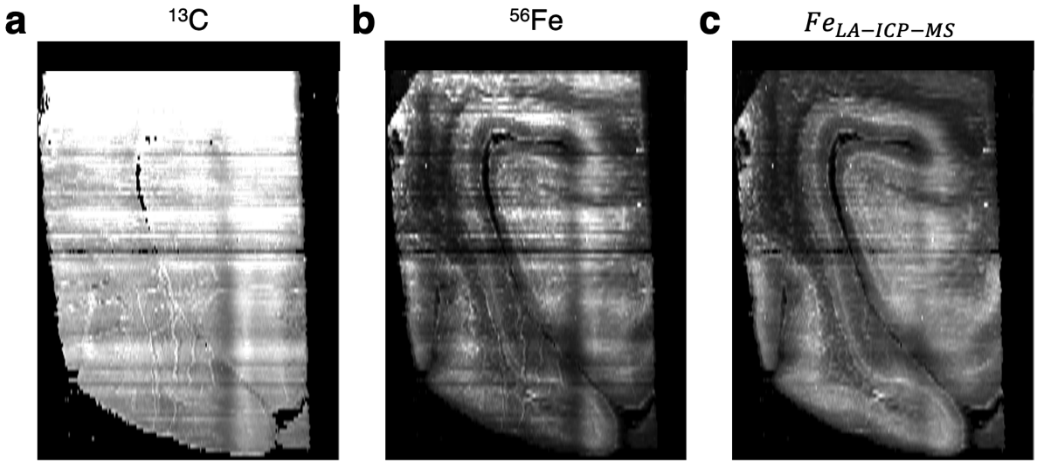


**Supplementary Figure 2. Original LA-ICP-MS images of (a) $^{13}C$ and (b) $^{56}Fe$, and (c) normalized $^{56}Fe$ image by $^{13}C$.** Because of the long scan time of the LA-ICP-MS imaging (~7 hours), the $^{13}C$ image was utilized to remove the signal drifts of LA-ICP-MS in the $^{56}Fe$ image. The normalization may affect iron quantification (Bulk et al., 2020).

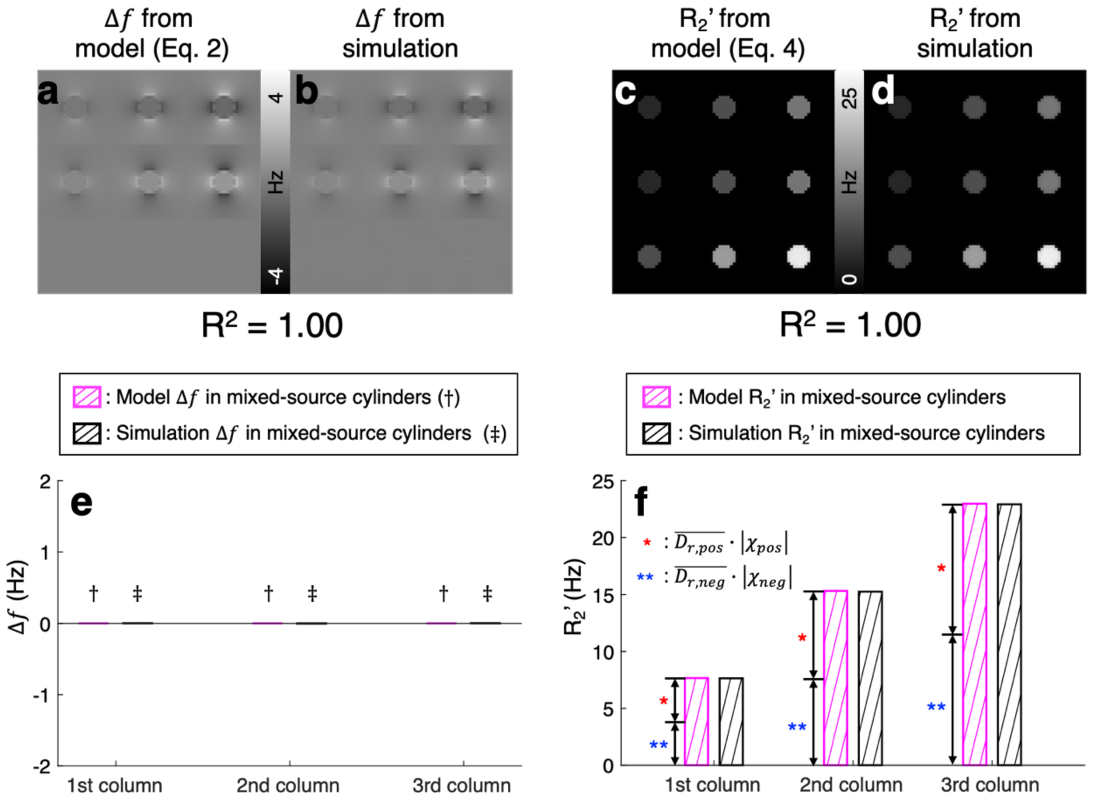


**Supplementary Figure 3. Validation of the proposed susceptibility models (Eqs. 2 and 4) using the Monte-Carlo simulation.** When the frequency shift maps from Eq. 2 are compared to those from the Monte-Carlo simulation, they reveal almost identical spatial distribution (**a-b**; $R^2 = 1.00$ in a voxel-wise linear regression). In particular, the simulation results of the mixed-source cylinders containing both positive and negative susceptibility sources (third row in **b**) report zero-frequency shifts, confirming our frequency shift model (Eq. 2), which states the signed sum of the positive and negative susceptibility determines the frequency shift. The quantitative frequency shift values of the mixed-source cylinders are shown in **e** with the pink and black bars representing the results from our model and the Monte-Carlo simulation, respectively (daggers and double daggers for the positions of the pink and black bars, respectively). In **c** and **d**, the $R_2'$ results are summarized, demonstrating a strong correlation between our $R_2'$ model in Eq. 4 and the simulation ($R^2 = 1.00$ in a voxel-wise linear regression). When the positive and negative susceptibility sources co-exist (third row in **d**), the $R_2'$ values from the simulation are determined by the (weighted) absolute sum of the positive and negative susceptibility, validating our $R_2'$ model in Eq. 4 (**f**). The single asterisk in **f** denotes the portion of the positive susceptibility in $R_2'$ from Eq. 4, whereas the double asterisk denotes that of the negative susceptibility.

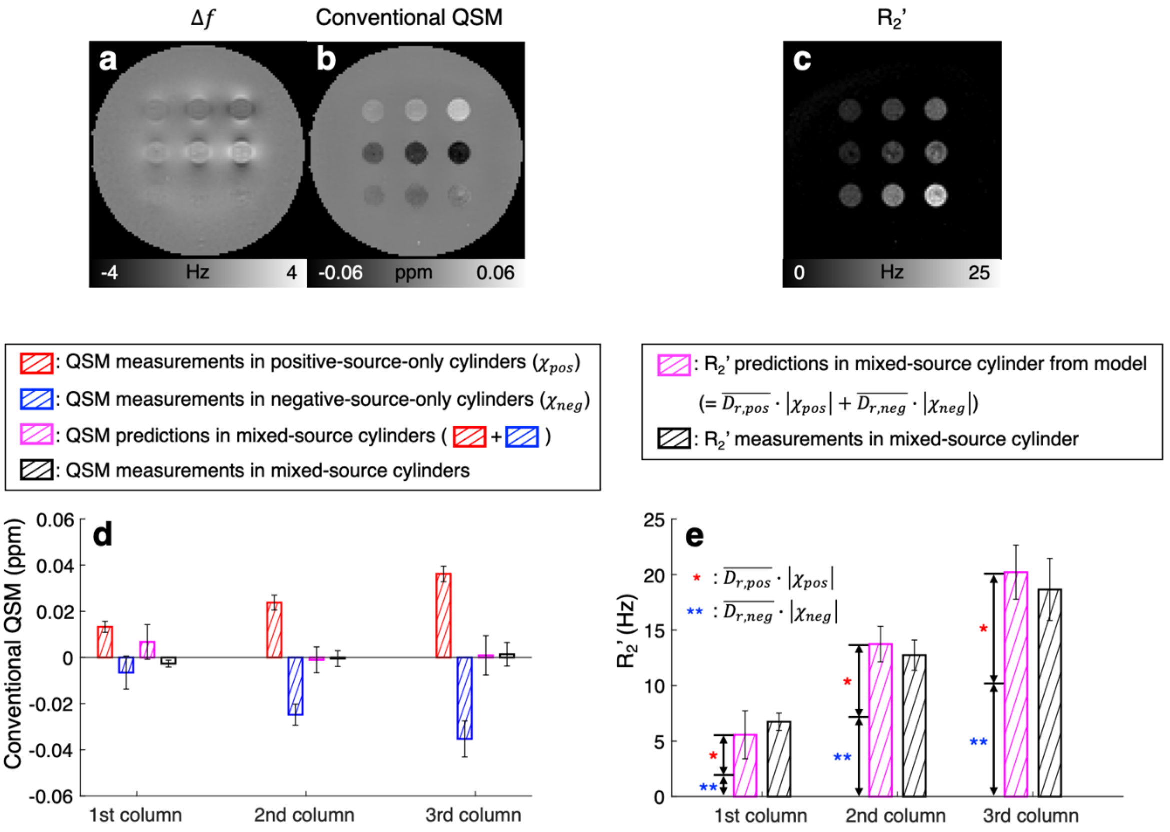


**Supplementary Figure 4. Validation of the proposed susceptibility models (Eqs. 2 and 4) using the phantom experiment. a**, Frequency shift map. **b**, QSM map. **c**, $R_2'$ maps. For quantitative analysis, the QSM results are utilized instead of the frequency shift results because of the non-local effects from adjacent cylinders and background field in the frequency shift map. The bar graph (**d**; mean ± standard deviation) reports the susceptibility concentrations of the positive-source-only cylinders (red bars) and the negative-source-only cylinders (blue bars). When the two measurements of each column are summed, they produced the QSM values in the pink bars. These measurements are close to the QSM values in the mixed-source cylinders (black bars) in all three columns. The results consolidate the validity of our model in Eq. 2. In the bar graph for $R_2'$ (**e**; mean ± standard deviation), the (weighted) absolute sums of the susceptibility measurements from the positive-source-only and negative-source-only cylinders (pink bars) match the $R_2'$ values in the mixed-source cylinders (black bars). The single and double asterisks in **e** represent the portions of the positive and negative susceptibility sources, respectively. The results corroborate our model in Eq. 4.

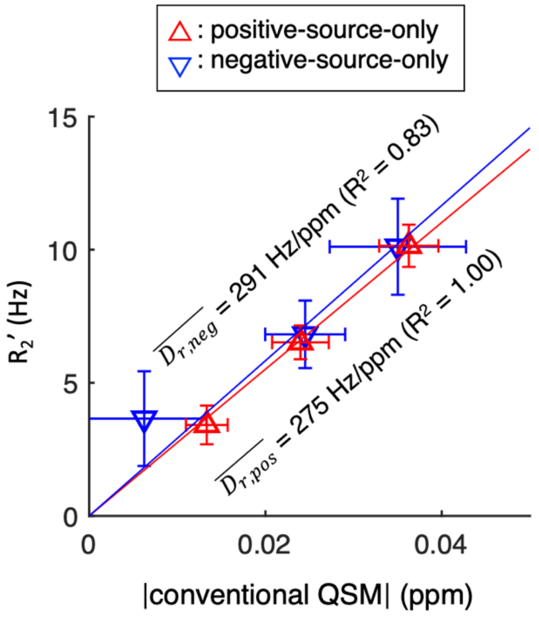


**Supplementary Figure 5. Estimation of the relaxometric constants in the phantom experiment.** The red upper triangles (or blue lower triangles) denote $R_2'$ values with respect to the absolute values of the conventional QSM in the positive-source-only (or negative-source-only) cylinders. Error bars indicate the standard deviations. When a linear regression was performed for the positive susceptibility source measurements, the slope ($\overline{D_{r,pos}}$) was 275 Hz/ppm ($R^2$ = 1.00). For the negative susceptibility source measurements, the slope ($\overline{D_{r,neg}}$) was 291 Hz/ppm ($R^2$ = 0.83).

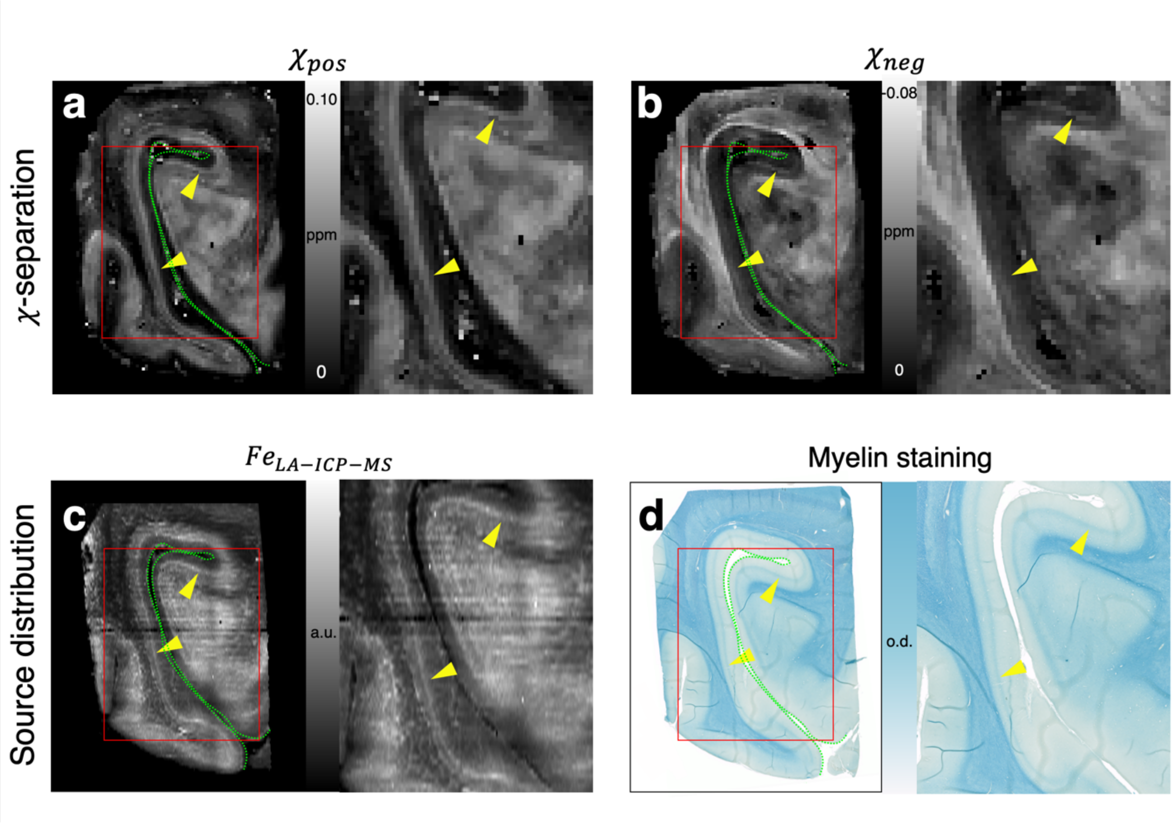


**Supplementary Figure 6. Zoomed-in images of Figure 4. a**, Positive susceptibility map. **b,** Negative susceptibility map. **c**, Iron image from LA-ICP-MS. **d**, Myelin image from LFB myelin staining. Red box areas, which contain the stria of Gennari (yellow arrowheads), are zoomed-in.

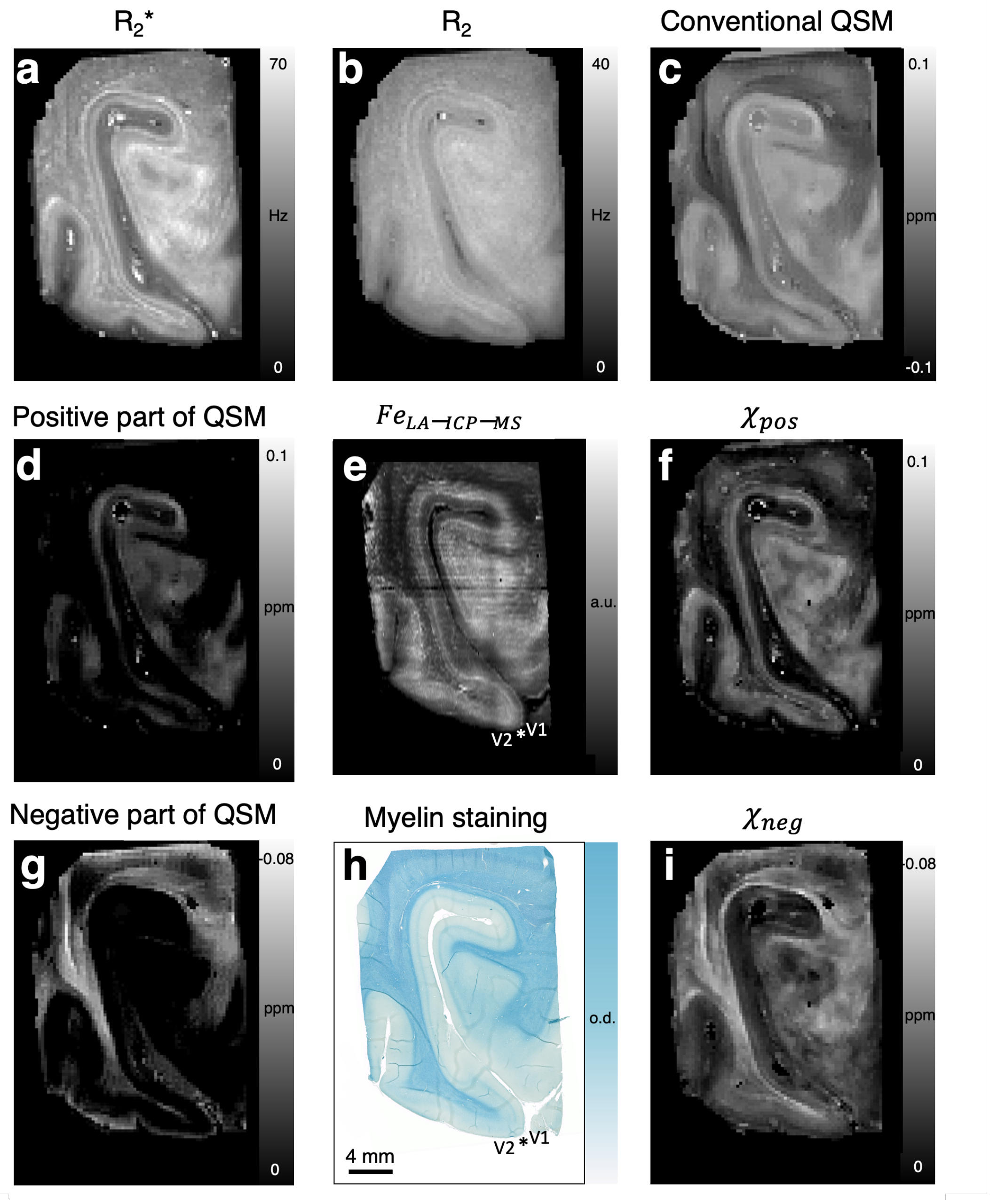


**Supplementary Figure 7. Comparison between MRI maps vs. iron and myelin images in the *ex-vivo* brain.** **a**, $R_2$* map. **b**, $R_2$ map. **c**, Conventional QSM map. **d**, Positive part of the conventional QSM map. **e,** Iron image from LA-ICP-MS. **f**, Positive susceptibility map. **g,** Negative part of the conventional QSM map. **h**, Myelin image from LFB myelin staining. **i,** Negative susceptibility map.

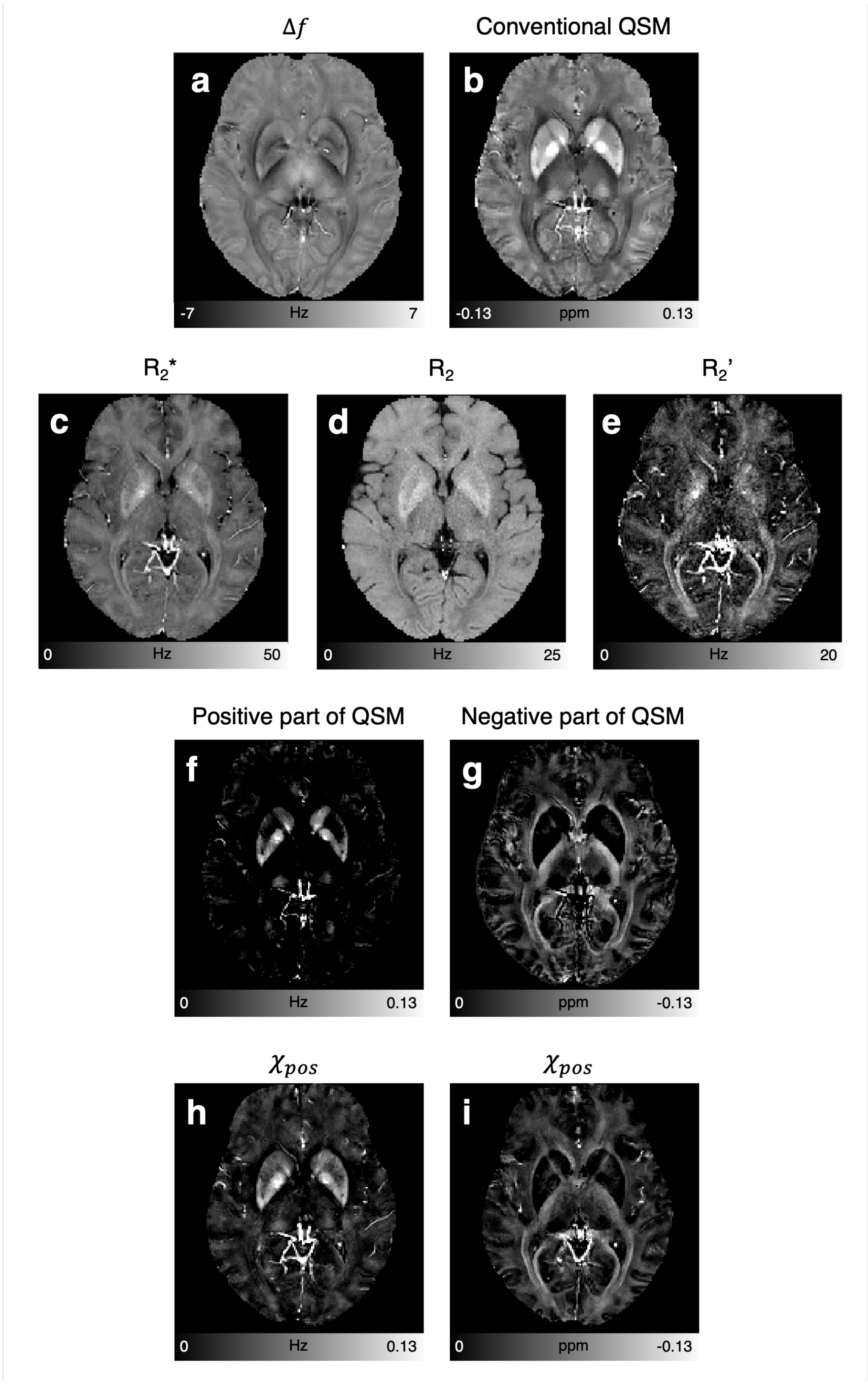


**Supplementary Figure 8. MRI maps in the *in-vivo* brain. a**, Frequency shift map. **b**, Conventional QSM map. **c**, $R_2$* map. **d**, $R_2$ map. **e**, $R_2$' map. **f**, Positive part of the conventional QSM map. **g**, Negative part of the conventional QSM map. **h**, Positive susceptibility map. **i**, Negative susceptibility map.

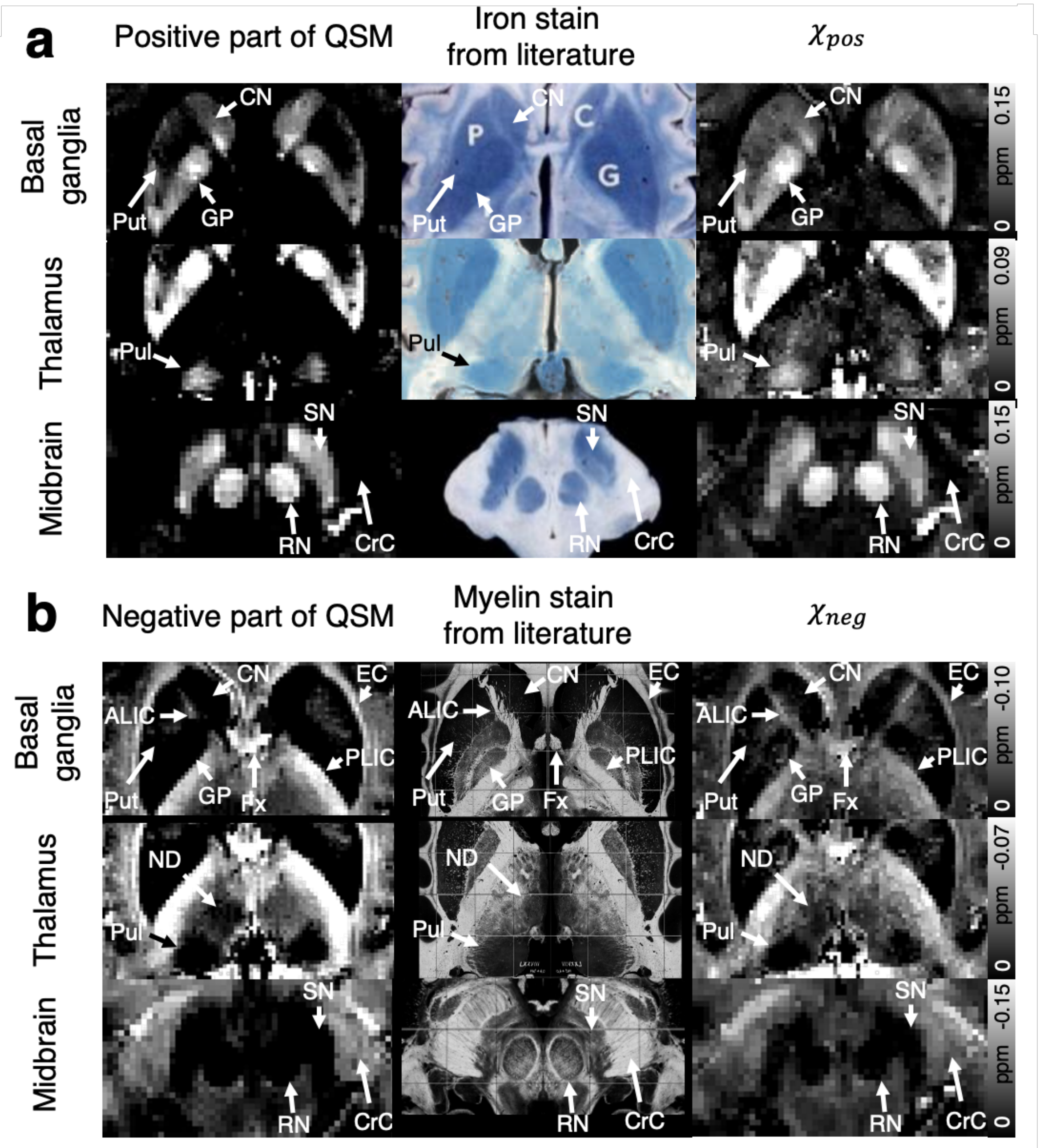


**Supplementary Figure 9. Comparison between the *in-vivo* MRI maps vs. iron and myelin histology from literatures. a,** Positive part of the conventional QSM maps (first column), iron staining images (second column), and positive susceptibility maps (third column). **b**, Negative part of the conventional QSM maps (first column), myelin staining images (second column), and negative susceptibility maps (third column).

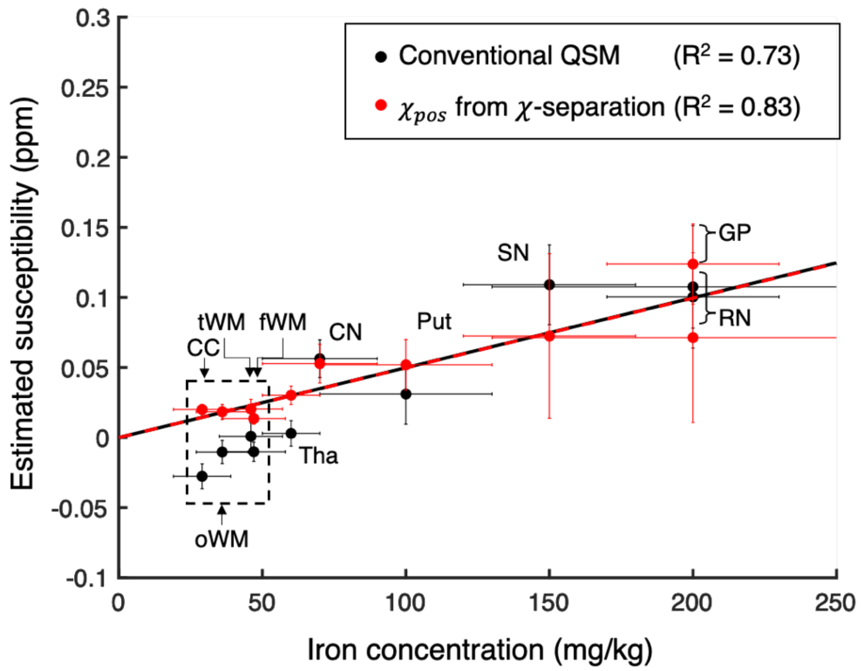


**Supplementary Figure 10. Positive susceptibility measurements from $\chi$-separation (red dots) and the susceptibility measurements from the conventional QSM (black dots) in ten ROIs:** thalamus (Tha), caudate nucleus (CN), putamen (Put), substantia nigra (SN), globus pallidus (GP), red nucleus (RN), and four regions in white matter (genu and splenium of corpus callosum (CC), occipital WM (oWM), temporal WM (tWM), and frontal WM (fWM); dashed box). When the ROI-averaged susceptibility measurements are plotted with respect to the literature values of iron concentrations (gray matter iron concentrations from Schweser et al. (Schweser et al., 2011); white matter iron concentrations from Langkammer et al. (Langkammer et al., 2012)), the positive susceptibility results from $\chi$-separation show a stronger linear relationship ($R^2 = 0.83$; red dashed line) than those from the conventional QSM ($R^2 = 0.73$; black solid line). Vertical error bars indicate the inter-subject standard deviations of the susceptibility values, whereas horizontal error bars denote the standard deviations of the iron concentrations from the literatures.

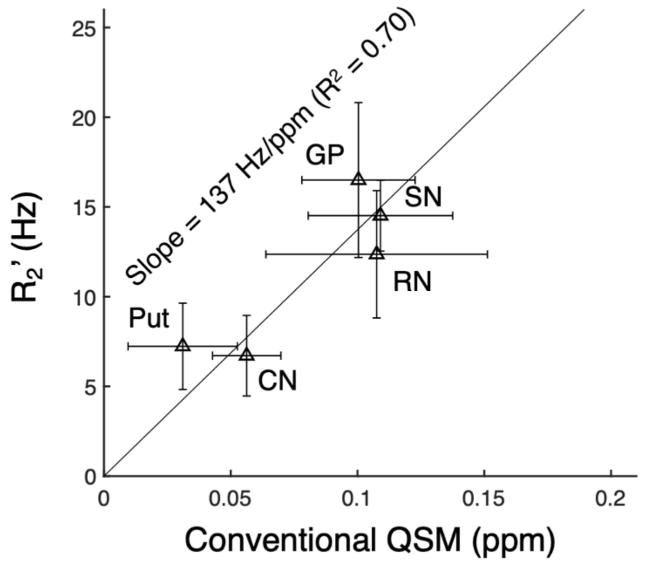


**Supplementary Figure 11. Estimation of the relaxometric constant in the *in-vivo* brains.** A linear regression between $R_2$' and conventional QSM values in the selected ROIs (Put, CN, RN, SN, and GP) reports the slope (i.e., $\overline{D_{r,pos}}$ ) of 137 Hz/ppm with $R^2$ of 0.70. Horizontal and vertical error bars indicate the inter-subject standard deviation of the susceptibility and $R_2$' values in each ROI.

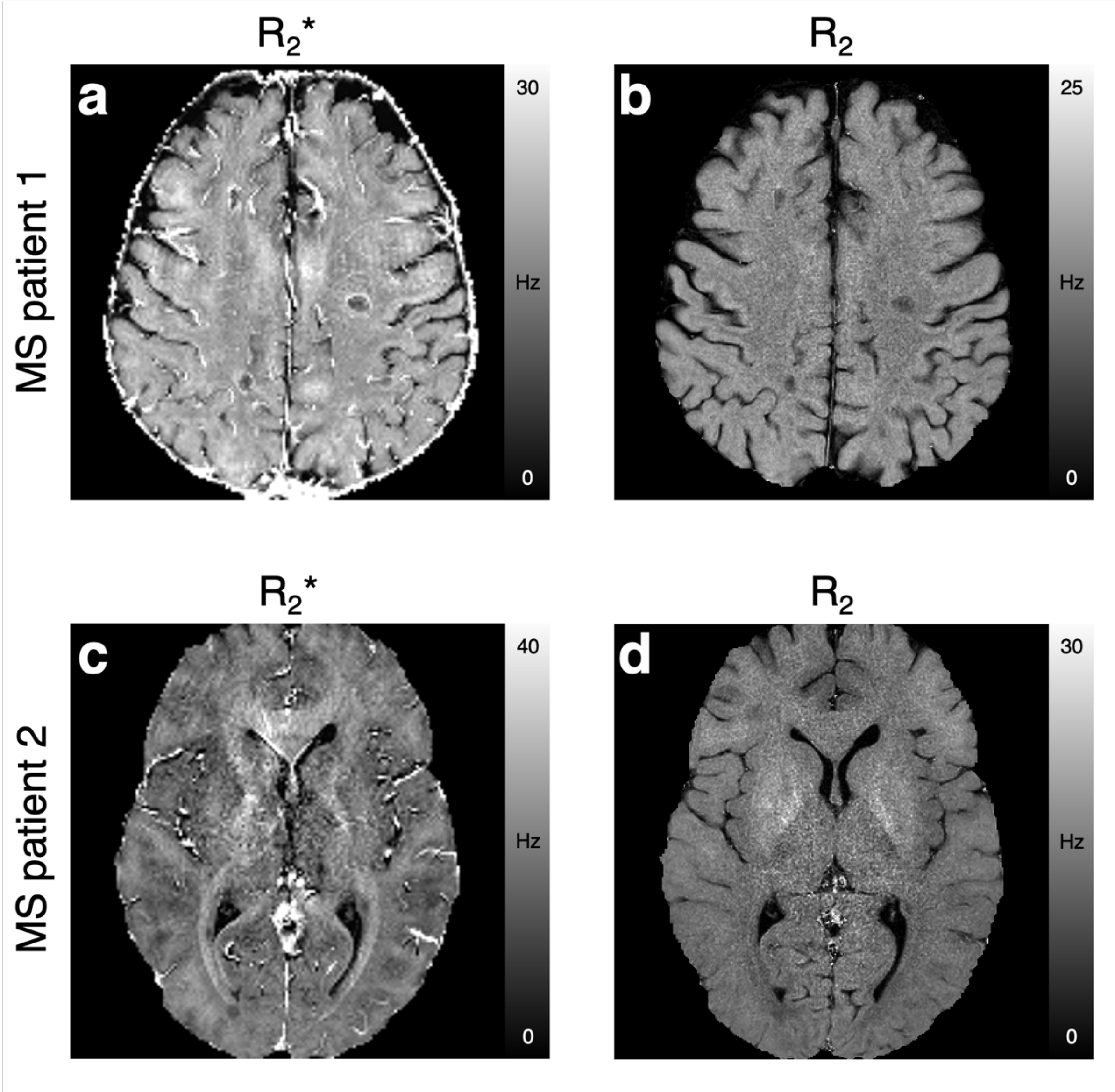


**Supplementary Figure 12. Additional *in-vivo* results in the two MS patients. a and c,** $R_2$* maps. **b and d,** $R_2$ maps.

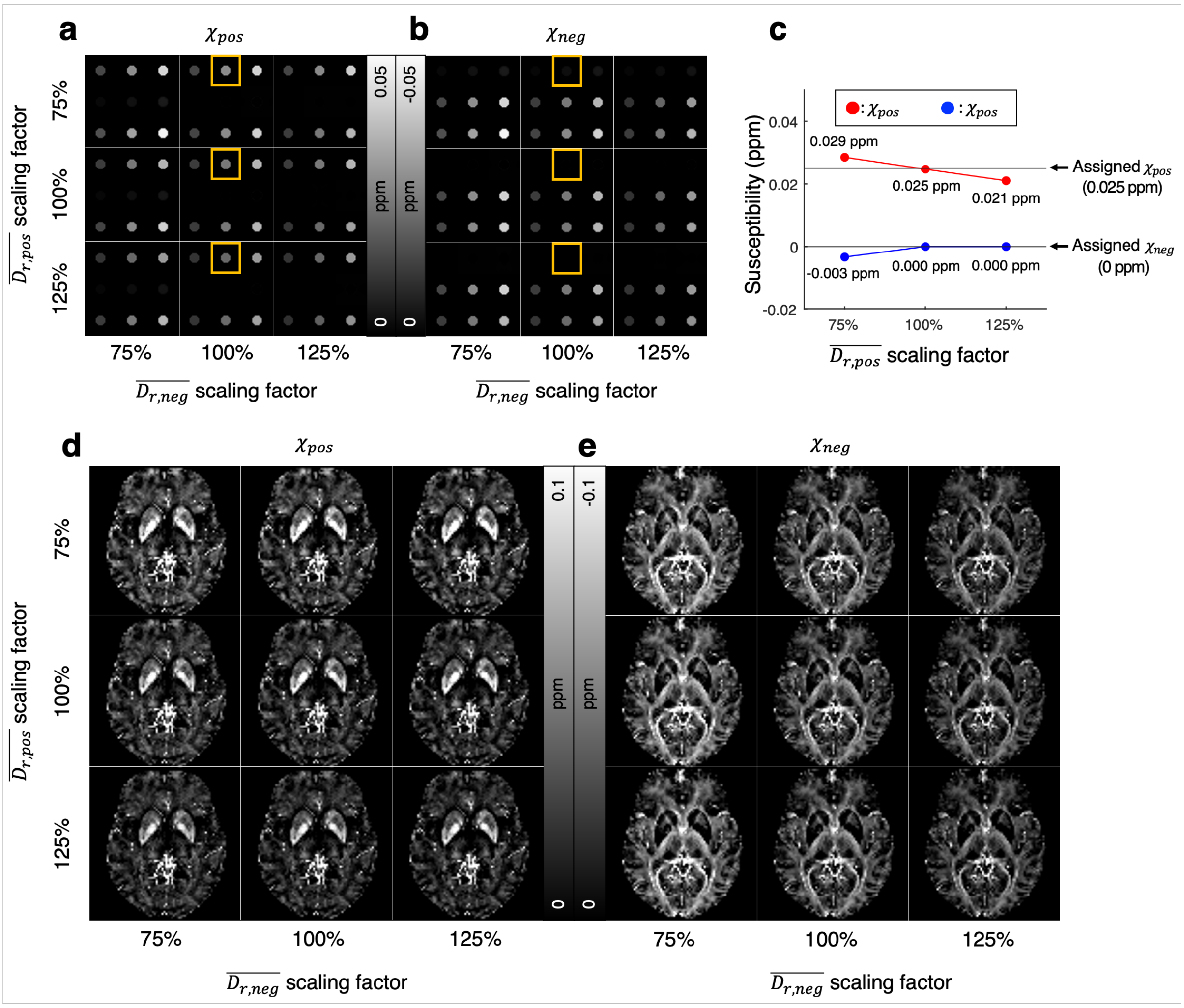


**Supplementary Figure 13. Effects of the relaxometric constants on (a-c) the Monte-Carlo simulation and (d-e) the *in-vivo* experiment.** When the positive and negative relaxometric constants were scaled between 75% to 125% of the original relaxometric constants, the positive and negative susceptibility maps show qualitatively similar contrasts, demonstrating the robustness of the $\chi$-separation method for the range of errors in the relaxometric constants (**a-b, d-e**). The quantitative results from the cylinders in the yellow boxes (**a-b**) reveal the relationship between relaxometric constants and susceptibility estimation errors (**c**).

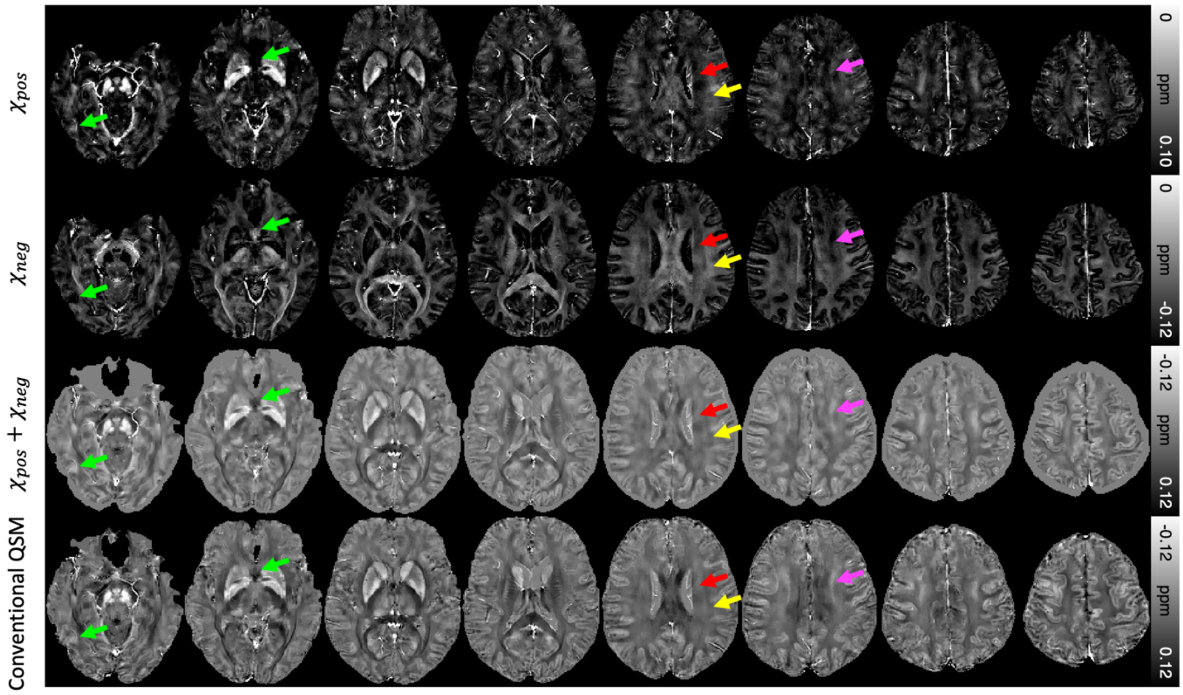


**Supplementary Figure 14. Representative slices of the *in-vivo* $\chi$-separation results.** First, second, and third rows display the positive susceptibility map, the negative susceptibility map, and the sum of the two maps, respectively. The last row shows the conventional QSM map. Green arrows denote artifacts from large $B_0$ field inhomogeneity. Red arrows indicate fibers of corona radiata that are largely parallel to the $B_0$ field, whereas yellow arrows point superior longitudinal fasciculus fibers that are approximately perpendicular to the $B_0$ field. The contrast difference between these two areas may be related not only to myelin concentration difference but also to fiber orientation-dependent anisotropic susceptibility (Lee et al., 2010; Wharton and Bowtell, 2012). Pink arrows indicate streaking artifacts induced from ventricular cerebrospinal fluid and deep gray matter, which have been observed in conventional QSM results (see coronal image from MEDI in Figure 2 of Yoon et al. (Yoon et al., 2018)).

**Supplementary note**

In this section, an analytical equation is derived to extend the linear relationship between $R_2$' and bulk susceptibility in Eq. 3 (Yablonskiy and Haacke, 1994) from susceptibility sources with a single source property (i.e., single characteristic frequency and geometry) to susceptibility sources with multiple source properties (i.e., *K* categories of the characteristic frequencies and/or geometries). The derivation is demonstrated for a spherical source geometry but can be extended for other geometries.

The notations for the following mathematical formula are listed below:

$V_0$: $volume\ of\ the\ medium$

$v_n$: $volume\ of\ the\ n^{th}\ susceptiblity\ source$

$v$: $volume\ of\ the\ total\ susceptibility\ sources$

$V$: $total\ volume\ including\ the\ sources\ and\ medium\ (= v + V_0)$

$\varsigma$: $volume\ fraction\ of\ the\ sources\ (= \frac{v}{V})$

$\varsigma_k$: $volume\ fraction\ of\ the\ sources\ in\ k^{th}\ category$

$\chi_0$: $susceptibility\ of\ the\ medium$

$\hat{\chi}_k$: $susceptibility\ of\ the\ source\ in\ k^{th}\ category$

$\overline{\chi_k}$: $bulk\ susceptibility\ of\ the\ source\ in\ k^{th}\ category\ relative\ to\ the\ medium$

$(= (\hat{\chi}_k - \chi_0) \cdot \varsigma_k)$

$N$: $total\ number\ of\ the\ sources$

$N_k$: $number\ of\ the\ sources\ in\ k^{th}\ category$

$B_0$: $applied\ magnetic\ field$

$M_0$: $magnetization\ density\ of\ the\ medium\ (= \chi_0 \cdot B_0)$

$M_k$: $magnetization\ density\ of\ the\ source\ in\ k^{th}\ category\ (= \hat{\chi}_k \cdot B_0)$

$b_n: magnetic\ field\ perturbed\ by\ n^{th}\ source$

$b_0: magnetic\ field\ in\ the\ medium\ in\ the\ absence\ of\ the\ source\ (= B_0 - \frac{8\pi}{3} \cdot M_0)$

$\omega(\vec{r}): frequency\ of\ the\ medium\ at\ point\ \vec{r}$

$\omega_n(\vec{r}): frequency\ of\ the\ medium\ at\ point\ \vec{r}\ generated\ by\ n^{th}\ source$

$\omega_0: reference\ frequency\ (= \gamma \cdot b_0)$

$\delta\omega_{s,k}: characteristic\ frequency\ of\ the\ spherical\ source\ in\ k^{th}\ category$

$(= \gamma \cdot \frac{4\pi}{3} \cdot (M_k - M_0))$

$R_n: radius\ of\ n^{th}\ source$

$\vartheta: angle\ relative\ to\ B_0$

$\rho: spin\ density\ of\ the\ medium$

$P_k(R): probability\ distribution\ of\ the\ radius\ of\ the\ source\ in\ k^{th}\ category$

*All units are in* CGS

In the presence of the susceptibility sources, the MRI signal from the medium is calculated as a volume integral of spin magnetizations, which experience frequency shifts from the susceptibility sources:

$$S(t) = \frac{1}{V} \cdot \rho \int_{V_0} d\vec{r} \cdot exp(-i \cdot \omega(\vec{r}) \cdot t) \qquad \text{[Eq. S1]}$$

where

$$\omega(\vec{r}) = \omega_0 + \sum_{n=1}^{N} \omega_n(\vec{r} - \vec{r_n}), \qquad \text{[Eq. S2]}$$

$$\omega_n(\vec{r} - \vec{r_n}) = \gamma \cdot b_n(\vec{r} - \vec{r_n}). \qquad \text{[Eq. S3]}$$

Since we assumed spherical susceptibility sources,

$$\omega_n(\vec{r}) = \delta\omega_{s,k_n} \cdot \left(\frac{R_n}{|\vec{r}|}\right)^3 (3cos^2\vartheta - 1) \quad \text{[Eq. S4]}$$

where $k_n$ denotes the category where the $n^{\text{th}}$ source is included ($k_n \in$ {1, 2, ..., K}).

When the sources are randomly distributed, the signal in Eq. S1 can be statistically averaged as follows:

$$\bar{S}(t) = \rho \cdot (1 - \varsigma) \cdot \prod_{n=1}^{N} \frac{1}{V - v_n} \cdot \int_{V - v_n} d\vec{r} \cdot exp(-i \cdot \omega_n(\vec{r}) \cdot t). \quad \text{[Eq. S5]}$$

In this equation, the field perturbation effects of the $N$ individual susceptibility sources are separated as a multiplication form. By substituting Eq. S4 into Eq. S5 and introducing the source radius $R$ as a random variable, Eq. S5 becomes

$$\bar{S}(t) = \rho \cdot (1 - \varsigma) \cdot \prod_{n=1}^{N} \int dR \cdot P_{k_n}(R) \cdot \frac{1}{V - v(R)} \cdot \int_{r>R} d\vec{r} \cdot exp\left(-i \cdot \delta\omega_{s,k_n} \cdot t \cdot \left(\frac{R}{r}\right)^3 (3cos^2\vartheta - 1)\right). \quad \text{[Eq. S6]}$$

When $N_k$ number of sources are included in the $k^{\text{th}}$ category (k ∈ {1, 2, ..., K}), Eq. S6 can be reformulated as follows:

$$\bar{S}(t) = \rho \cdot (1 - \varsigma) \cdot \prod_{k=1}^{K} \left\{\int dR \cdot P_k(R) \cdot \frac{1}{V - v(R)} \cdot \int_{r>R} d\vec{r} \cdot exp\left(-i \cdot \delta\omega_{s,k} \cdot t \cdot \left(\frac{R}{r}\right)^3 (3cos^2\vartheta - 1)\right)\right\}^{N_k} \quad \text{[Eq. S7]}$$

This equation can be simplified as follows (see Appendix in Yablonskiy and Haacke (Yablonskiy and Haacke, 1994)):

$$|\bar{S}(t)| = \rho \cdot (1 - \varsigma) \cdot \prod_{k=1}^{K} \left(1 - \frac{\varsigma_k}{N_k} \cdot \frac{2 \cdot \pi}{3 \cdot \sqrt{3}} \cdot \left|\delta\omega_{s,k}\right| \cdot t\right)^{N_k} \quad \text{[Eq. S8]}$$

assuming $\left|\delta\omega_{s,k}\right| \cdot t \gg 1$ for all $k$. This assumption is valid for ferritin particles at 3 T ($\delta\omega_s$ > $10^5$ Hz) for a typical echo time (> 1 ms).

Considering a large number of sources for all categories (i.e., $N_k \to \infty$ for all $k$), Eq. S8 is simplified as an exponential function:

$$|\bar{S}(t)| = \rho \cdot (1-\varsigma) \cdot exp\left[-\frac{2\cdot\pi}{3\cdot\sqrt{3}} \cdot \left(\sum_{k=1}^{K}\left|\delta\omega_{s,k}\right| \cdot \varsigma_k\right) \cdot t\right]. \qquad \text{[Eq. S9]}$$

Using the definition, we can substitute $\left|\delta\omega_{s,k}\right| \cdot \varsigma_k$ as $\gamma \cdot \frac{4}{3} \cdot \pi \cdot |\hat{\chi}_k - \chi_0| \cdot B_0 \cdot \varsigma_k = \frac{4}{3} \cdot \pi \cdot \gamma \cdot B_0 \cdot |\overline{\chi_k}|$. Finally, the exponential decay constant in Eq. S9, which is equivalent to a $R_2$' relaxation rate, is represented as follows:

$$\sum_{k=1}^{K} \frac{2\cdot\pi}{9\cdot\sqrt{3}} \cdot \gamma \cdot B_0 \cdot |4 \cdot \pi \cdot \overline{\chi_k}| = \sum_{k=1}^{K} \frac{2\cdot\pi}{9\cdot\sqrt{3}} \cdot \gamma \cdot B_0 \cdot \left|\overline{\chi_{k,SI}}\right| = \sum_{k=1}^{K} D_{r,k} \cdot \left|\overline{\chi_{k,SI}}\right| = R_2'.$$

[Eq. S10]

where $\overline{\chi_{k,SI}}$ is equivalent to the bulk susceptibility value in the SI unit and $D_{r,k}$ is the relaxometric constant that describes the contribution of the bulk susceptibility of the $k^{th}$ category source to $R_2$'. This result demonstrates that $R_2$' is proportional to the absolute sum of the bulk susceptibility values of the individual category.

For two categories of the susceptibility sources (i.e., K = 2) with the opposite sign (i.e., $\delta\omega_{s,1} > 0$, and $\delta\omega_{s,2} < 0$), Eq. S10 is simplified to Eq. 4, explaining the effects of positive and negative sources on $R_2$'. The relaxometric constant can be affected by water diffusion, and the size, geometry, and characteristic frequency of the susceptibility sources (Yung, 2003).
